# Multi-omic longitudinal profiling of companion dogs reveals cross-omic coordination and structured aging trajectories

**DOI:** 10.64898/2026.11.12.764519

**Authors:** Tal Bamberger, Yadid M. Algavi, Efrat Muller, Omri Peleg, Shiri Baum, Ashlee Greenier, Christine Adjangba, Elizabeth Slikas, Layla Brassington, Blaise Mariner, Brianah McCoy, Benjamin R. Harrison, Maria Partida-Aguilar, Abbey Marye, Adam Harris, Emily Rout, DAP Consortium, Anne Avery, Daniel E.L. Promislow, Noah Snyder-Mackler, Elhanan Borenstein

## Abstract

Aging is a systems-level biological process shaped by coordinated changes across molecular, physiological, and environmental domains, yet how these domains interact and impact aging remains poorly understood. Companion dogs offer a powerful model for addressing this gap, combining shared human environments with diversity in genetics, lifespan, and disease susceptibility. Here, we present a longitudinal multi-omic analysis of aging in 965 companion dogs from the Dog Aging Project, integrating microbiome, epigenome, blood metabolome, and immune phenotyping with extensive clinical, environmental, and lifestyle surveys. We reveal an interconnected cross-omic architecture, with age emerging as the dominant axis of biological variation. We further identify a predictive multi-omic aging signature, distinguish linear from non-linear aging trajectories, uncover breed size-dependent dynamics, and characterize omic-specific mortality risk. Collectively, our findings reveal organizing principles of aging, many paralleling human observations, provide a unique resource for integrative geroscience, and establish dogs as a translational model for aging research.

## Introduction

Aging is a universal biological process and the dominant risk factor for most chronic diseases, functional decline, and mortality^1–3^. Despite its universal nature, individuals differ widely in the rate, timing, and ordering of physiological deterioration and disease onset. This heterogeneity underscores aging as a systems-level phenomenon that is shaped by complex interactions between an individual’s genetic background, molecular regulation, environmental exposures, and lifestyle^4,5^. Understanding how these interacting processes change over time, and how their trajectories impact aging and survival, remains a central challenge in aging biology.

To date, much of our understanding of the molecular mechanisms involved in aging has been gained via controlled experimental studies in laboratory mammalian model organisms^6,7^. Research in captive mice and primates, for example, has identified conserved molecular pathways and hallmarks of aging and enabled causal interrogation of aging through genetic and environmental perturbations^8–11^. However, such insights were derived largely from genetically homogeneous populations maintained under highly controlled conditions that minimize environmental variation, pathogen exposure, and lifestyle diversity^12,13^. As a result, the extent to which mechanisms identified in laboratory settings capture the determinants of aging in outbred populations living in complex, real-world environments remains uncertain.

In parallel, observational studies of aging in natural settings have expanded rapidly, encompassing both population-wide human cohorts and longitudinal studies of free-living animal populations, including studies in squirrels^14^, meerkats^15^, horses^16^, primates^17–19^, and other mammals^20^. These studies capture aging in genetically diverse populations exposed to ecologically-and environmentally-relevant conditions, revealing substantial inter-individual variation in aging trajectories. Yet, from a translational point of view, insights obtained from observational studies of aging in free-living animal populations may not carry over to human populations. In industrialized societies, human aging is profoundly shaped by medical intervention^21^, stable and abundant nutrition^22,23^, and extensive technological environmental modification^24^. By contrast, animals in natural ecosystems experience fundamentally different dietary regimes, disease pressures, physical activity patterns, and ecological constraints^25^. Consequently, molecular signatures of aging observed in humans and in wild animal populations may reflect distinct environmental contexts as much as conserved biological processes, complicating efforts to integrate findings across species and to bridge experimental and observational aging research.

Companion dogs occupy a unique position that helps to resolve this comparability gap. Living alongside humans, dogs share much of the built environment, diet composition, healthcare access, and lifestyle-associated exposures characteristic of industrialized societies with their human owners^26^. Importantly, dogs also develop many of the same age-associated diseases as humans^27^. At the same time, dogs are genetically diverse and exhibit substantial variation in body size, lifespan, and aging rate^28^. Crucially, dogs age approximately six to ten times faster than humans, enabling longitudinal investigation of aging processes and survival outcomes over experimentally tractable timescales. Combined, these features position companion dogs as a powerful model for studying aging in a shared environmental context across species.

Here, we present a comprehensive multi-omic analysis of aging in the Dog Aging Project (DAP) Precision Cohort–a large, open-science, longitudinal study of companion dogs^29,30^. We integrate gut microbiome profiling, DNA methylation, blood metabolomics, and immune cell phenotyping by flow cytometry with detailed phenotypic, clinical, and environmental exposure data across 965 companion dogs. We map how distinct molecular systems age relative to one another, quantify their coordination and temporal dynamics, and characterize the relationship between the various omics, aging, and mortality risk. Furthermore, we contrast identified canine aging dynamics with those observed in human multi-omic cohorts, highlighting conserved organizing principles. Collectively, our study provides an unprecedented, multi-omic, and high-resolution atlas of systems-level aging in a real-world, translationally meaningful environment, advancing our understanding of biological aging and serving as a foundational resource for future geroscience research.

## Results

### The Dog Aging Project (DAP) Precision Cohort

In this study, we analyzed longitudinal multi-omic data from the Precision Cohort of the Dog Aging Project, comprising 965 companion dogs profiled annually over two years (Fig. 1A). The cohort spanned a broad range of ages upon enrolment (mean = 6.09 years, std = 2.93; Fig. 1A, upper inset), and captured broad breed heterogeneity, including mixed-breed and purebred dogs (n = 451 and 514, respectively; Fig. 1B). Geographically, the cohort represents a demographically diverse population sampled across the United States (Fig. 1C). It included 492 males and 473 females, of whom 79.7% and 84.8% were sterilized, respectively (Fig. 1D). The cohort also spanned the full range of American Veterinary Medical Association (AVMA) breed-size classes and life stages (Fig. 1E). Owner-reported health status indicated that most dogs were in good health at enrollment (Fig. 1F).

**Figure 1:**
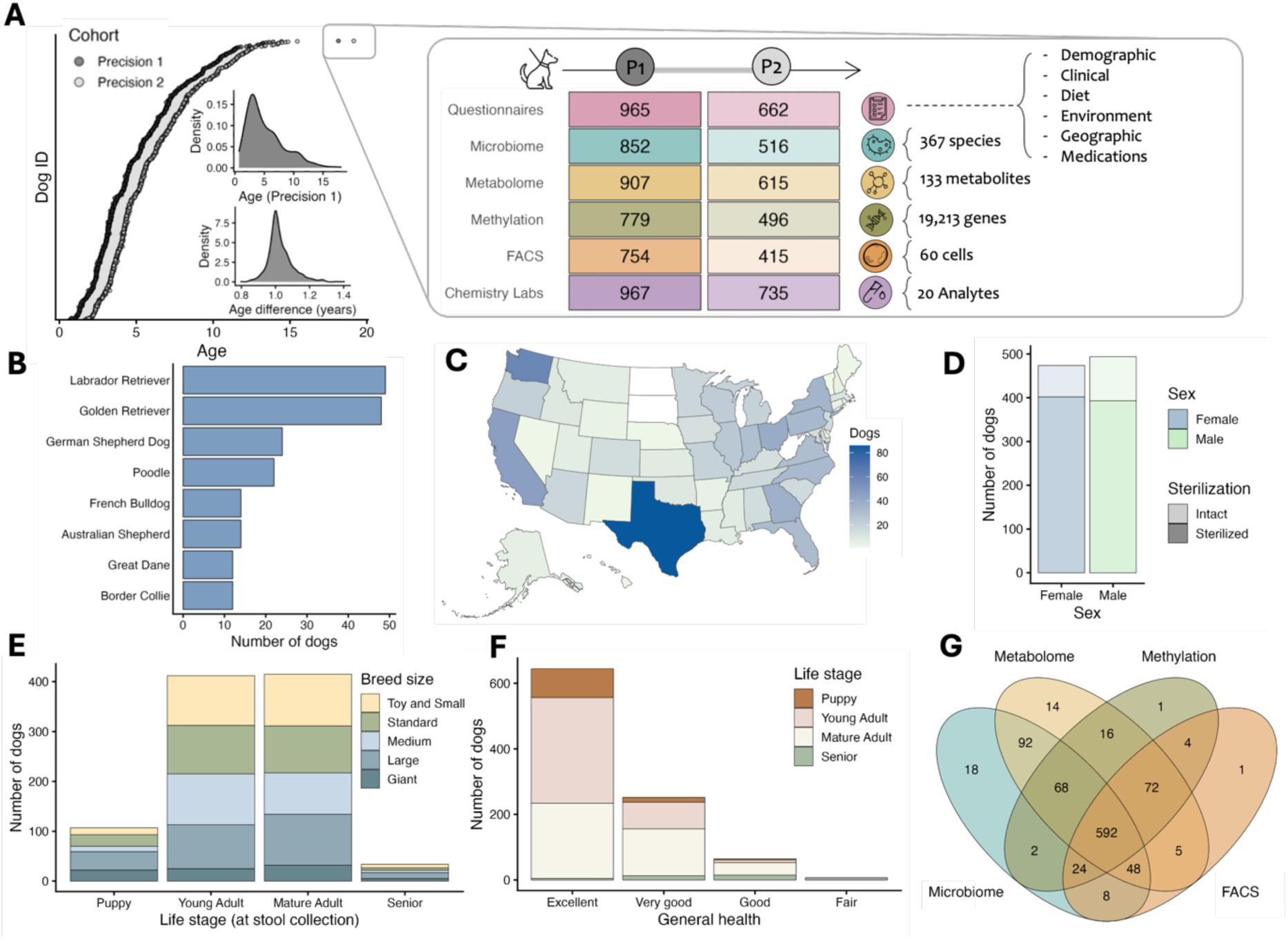
Characteristics of the DAP precision cohort. **(A)** Longitudinal sampling across two time points (Precision 1 and Precision 2). Right: each line represents one dog, with points marking the dog’s age at the two sampling times. Upper inset: age distribution in Precision 1. Lower inset: within-dog age differences. Left: a zoom-in illustrating available data for each dog with its Precision 1 and Precision 2 samples and data modalities (questionnaires, gut microbiome, metabolome, DNA methylation, flow cytometry, clinical chemistry). The accompanying table summarizes sample availability by modality and time point; cell hue encodes modality and shading encodes sample counts, with text detailing measured features per domain. **(B–G)** Precision 1 cohort characteristics: **(B)** Breed composition showing the eight most common purebred breeds (with all others aggregated into the “Other” category). **(C)** Map of the geographic distribution of enrolled dogs across the United States. **(D)** Stacked bar plot of sex, colored by sterilization status. **(E)** Stacked bar plot of breed size by life stage (AVMA categories, with colors denoting breed-size classes). **(F)** Stacked bar plot of owner-reported general health by life stage, with colors denoting life stage. **(G)** A Venn diagram of the various omics’ coverage and intersections in Precision 1. The numbers listed in the intersections denote the number of dogs with matched data across multiple omics, and thus available for integrative multi-omic analyses.

All 965 dogs in this cohort were sampled in the first year (Precision 1 [P1]), of which 662 dogs were resampled in the second year (Precision 2 [P2]; Fig. 1A), with a mean interval between sampling of 1.03 years (std = 0.07; Fig. 1A, lower inset). At each time point, dogs were extensively profiled across multiple omic domains. Gut microbiome composition was characterized by metagenomic shotgun sequencing of stool samples, resulting in composition profiles detailing the relative abundances of 367 microbial species^31^. The plasma metabolome was assessed using targeted LC-MS, quantifying 133 aqueous metabolites^32^. Epigenetic profiles were generated using reduced representation bisulfite sequencing (RRBS), providing DNA methylation measurements at 19,213 promoter-proximal CpG sites in peripheral blood mononuclear cells (PBMCs)^33,34^. Immunophenotyping was performed by flow cytometry, resulting in 60 features representing major leukocyte populations and their surface marker expression. Finally, all dogs underwent a veterinary assessment, resulting in extensive longitudinal phenotypic data including signalment, clinical, dietary, environmental, geographic, and medication information. While some omic profiles were missing for subsets of dogs, 592 dogs had complete profiles available across all domains in the first wave of the Precision cohort sampling (Precision 1; Fig. 1G). In Precision 2, fewer samples were available per omic, with 275 dogs retaining full multi-omic coverage.

### Global cross-omic concordance highlights microbiome-metabolome coupling

We first assessed global cross-omic associations in the Precision 1 cohort using sparse canonical correlation analysis (sCCA) to compare the shared variation between samples across omics. This analysis showed that all omic pairs are significantly associated (FDR < 0.1; Fig. 2A). The strongest alignment was observed between methylation and FACS profiles (shared variance, *r*^2^ = 29.7%), followed by the concordance between the microbiome and metabolome (*r*^2^ = 23.3%). These strong associations are perhaps not surprising, as methylation and FACS were both measured from the same type of biospecimen (namely, neutrophil-enriched PBMCs), and epigenetic patterns and immune-cell composition are also tightly interconnected through coordinated regulation of immune activation^35^. Similarly the gut microbiome is closely linked to host metabolic profiles^36^. As an additional validation, we performed Procrustes analysis to compare the structure of pairwise sample distances across omics, which revealed similar global trends (Supplementary Fig. 1).

**Figure 2.**
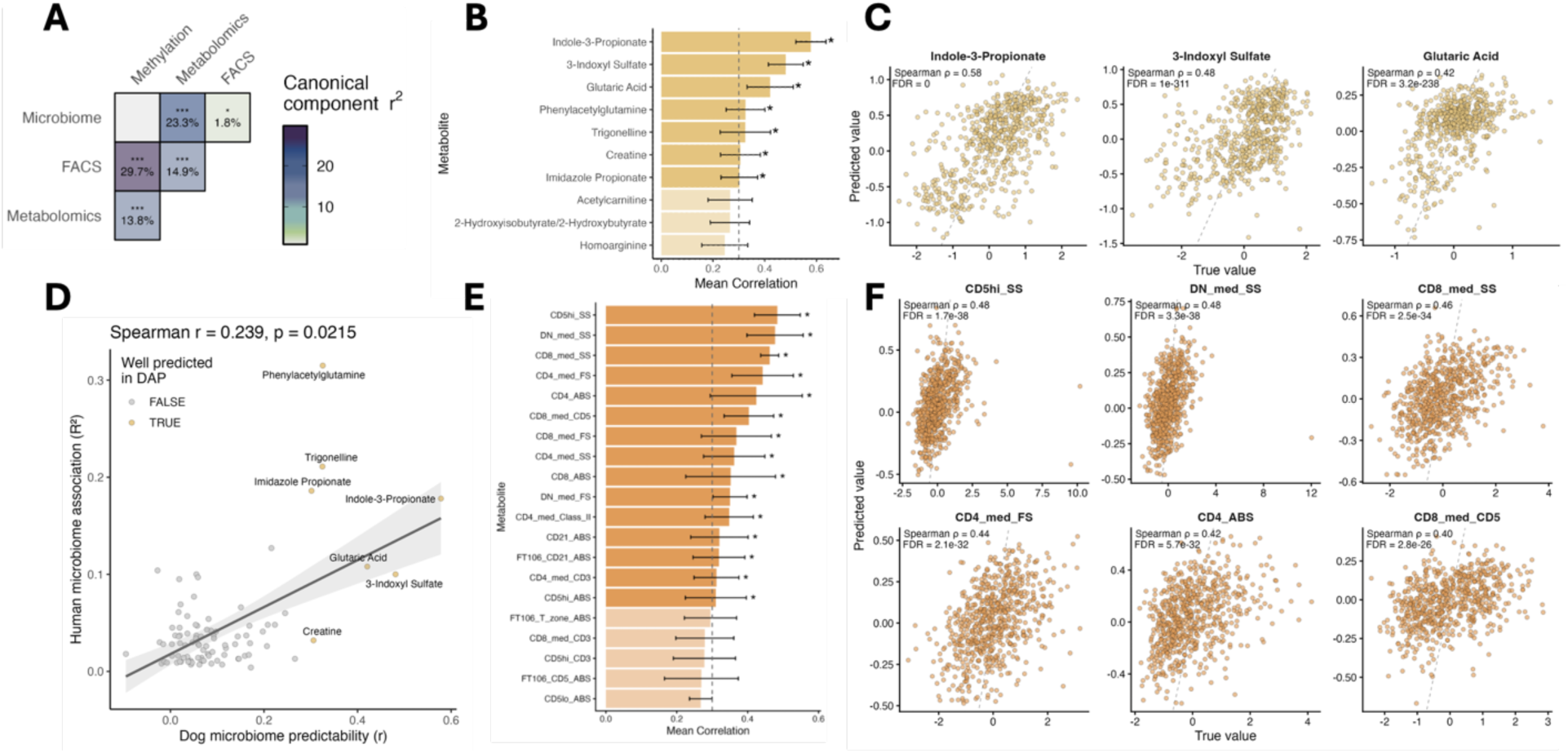
Cross-omic and microbiome-metabolome associations. **(A)** Sparse canonical correlation analysis quantifying shared variation between omic profiles; tiles are colored by the canonical component r^2^ (higher = stronger concordance). Asterisks denote FDR-adjusted significance (· FDR ≤ 0.1; * ≤ 0.05; ** ≤ 0.01; *** ≤ 0.001). **(B)** Predictability of metabolites: top 10 metabolites ranked by cross-validated correlation (Absolute Spearman ρ) between observed and predicted concentrations. Well predicted metabolites (Absolute Spearman ρ between real and predicted values, FDR < 0.1) are marked with an asterisk and shown in saturated color (and see complete results in Supplementary Fig. 2A and Table 1). **(C)** Observed vs. predicted values for the top three metabolites (held-out folds); all seven well-predicted metabolites are shown in Supplementary Fig. 2B. **(D)** Scatter plot comparing microbiome-based metabolite predictability in our DAP cohort (x-axis) with the variance explained by the microbiome in the SCAPIS human serum cohort^37^ (y-axis). Each point represents a shared metabolite. Metabolites classified as well predicted in the DAP cohort (Spearman r > 0.3, FDR < 0.1) are highlighted in yellow. **(E)** Predictability of FACS-derived immune-cell features from methylation profiles, shown as in panel B, with the top 20 FACS features. **(F)** Observed vs. predicted values for the top six FACS-derived immune-cell features, shown as in panel C; all fifteen well-predicted FACS features are shown in Supplementary Fig. 3B

To further dissect the strongest cross-omic relationships, we separately examined the two strongest cross-omic associations: microbiome–metabolome and methylation–immune system. We first sought to identify specific circulating metabolites that can be predicted from gut microbial composition. Using microbiome-based predictive models, we identified seven metabolites that were successfully predicted from species-level profiles (Spearman’s r > 0.3; FDR < 0.1; Fig. 2B,C; Supplementary Fig. 2), including microbiota-derived metabolites with established links to host physiology, such as indole-3-propionate, 3-indoxyl sulfate, and phenylacetylglutamine. Feature-importance analysis further indicated that these predictions were driven by coordinated contributions from multiple microbial taxa rather than by single dominant species (Supplementary Fig. 2C). Moreover, comparison with a large human cohort revealed that metabolites most predictable from the microbiome in dogs also tended to exhibit stronger microbiome dependence in humans, suggesting partial conservation of microbiome–metabolome interactions across species (Fig. 2D). See Supplementary Text and Supplementary Tables 1-3 for full results and interpretation.

Similarly, given the strong global association between methylation and FACS profiles, we examined whether PBMC methylation profiles capture variation in immune-cell composition. These analyses identified multiple immune-cell features that could be accurately predicted from methylation profiles, further supporting tight coupling between the epigenome and the immune system (Supplementary Text; Fig. 2E,F; Supplementary Fig. 3; Supplementary Tables 4,5).

### Longitudinal stability of omic-specific and cross-omic profiles

We next turned to characterize longitudinal dynamics and stability within and across omics. We specifically focused on dogs sampled at both time points (Precision 1 and Precision 2), and quantified feature-wise stability as the Spearman correlation between P1 and P2 for each feature in each omic separately across dogs (Supplementary Fig. 4A,B). Metabolome features showed the highest temporal stability (r = 0.444 ± 0.175, mean feature-wise Spearman correlation across 133 metabolites), followed by FACS (0.439 ± 0.230, across 60 FACS-derived features), microbiome (0.373 ± 0.086, 367 species), and methylation (0.069 ± 0.120, 19,213 sites) features. Consistent with these findings, global cross-time concordance (assessed via Procrustes tests between sample-wise distances at P1 and P2) was significant for all omics and showed a somewhat similar ordering (with Procrustes statistics of 19.6%, 17.5%, 17.1%, and 3.2% for FACS, microbiome, metabolome, and methylation, respectively; FDR-adjusted p < 0.1; Supplementary Fig. 4C).

We further found that across all omic profiles, dog-specific distances in multidimensional feature space between each dog’s P1 and P2 profiles were consistently and significantly smaller than the mean distance between that dog’s P1 profile and the P2 profiles of all other dogs (Supplementary Fig. 4D; FDR < 0.1, linear mixed-effects model adjusting for breed size, sex, and life stage; see Methods). Distances were calculated using Bray-Curtis dissimilarity for the microbiome and Euclidean distance for the metabolome, methylation, and FACS profiles. This observation supports overall stability in dogs’ molecular status and strong subject-specific trajectories. Moreover, ranking factors from most to least stable, we found that dogs that were older or less healthy tended to have larger longitudinal shifts averaged across omics (Supplementary Fig. 4E). Similarly, sex and sterilization status were associated with greater change: intact males showed larger P1 to P2 distances than neutered males and spayed females in the metabolome and methylation, suggesting a hormonal component to longitudinal variability.

To further examine whether *shared* multi-omic structure was similarly stable over time, we applied MOFA+^38^, a probabilistic multi-view factor model that reconstructs a low-dimensional latent representation capturing variation shared across omics. Applying MOFA+ to the four omics in our data, we found that the learned latent factors explained substantial variance within the different omic domains, with many factors driven primarily by individual domains and a subset showing contributions across multiple domains (Supplementary Fig. 5A,B). Consistent with our prior analyses, the MOFA+ latent factors were longitudinally stable (r = 0.370 ± 0.130, mean feature-wise Spearman correlation across 15 factors) and exhibited significant multivariate concordance between P1 and P2 in the MOFA+ latent space (Procrustes statistic = 15.3%, FDR-adjusted p < 0.1; Supplementary Fig. 4C). Similarly, dog-specific P1 to P2 distances in the MOFA+ latent space were significantly smaller than the corresponding mean distances to other dogs, based on the same linear mixed-effects model used above (FDR < 0.1; Supplementary Fig. 4D).

### The landscape of phenotype-omic associations highlights age as a central axis

To systematically map the potential impact of phenotypic and environmental factors on canine biology, we next quantified the association between these factors and molecular features. Phenotypic information in our dataset included comprehensive owner-reported survey data, spanning signalment, behavior, health conditions, dietary habits, environmental and geographic exposures, and medications. We hypothesized that while some traits might act as global modulators of canine biology, with observed associations across multiple omics, others would have more localized, omic-specific effects. Indeed, applying PERMANOVA analysis to each omic’s distance matrix and with each phenotype as a group label revealed a strikingly non-uniform association landscape (Fig. 3A, Supplementary Table 6). Specifically, signalment traits, particularly age and breed size, were consistently associated with molecular variation in every omic domain, suggesting that these traits represent key physiological axes that are linked to molecular processes in dogs. In contrast, the association of lifestyle and environmental factors with the various omic profiles was more granular. For instance, dietary components and some environmental exposures (including ozone or geographic region) were associated with variation in the microbiome and blood chemistry, respectively, rather than with broader, systemic shifts across multiple omic domains.

**Figure 3.**
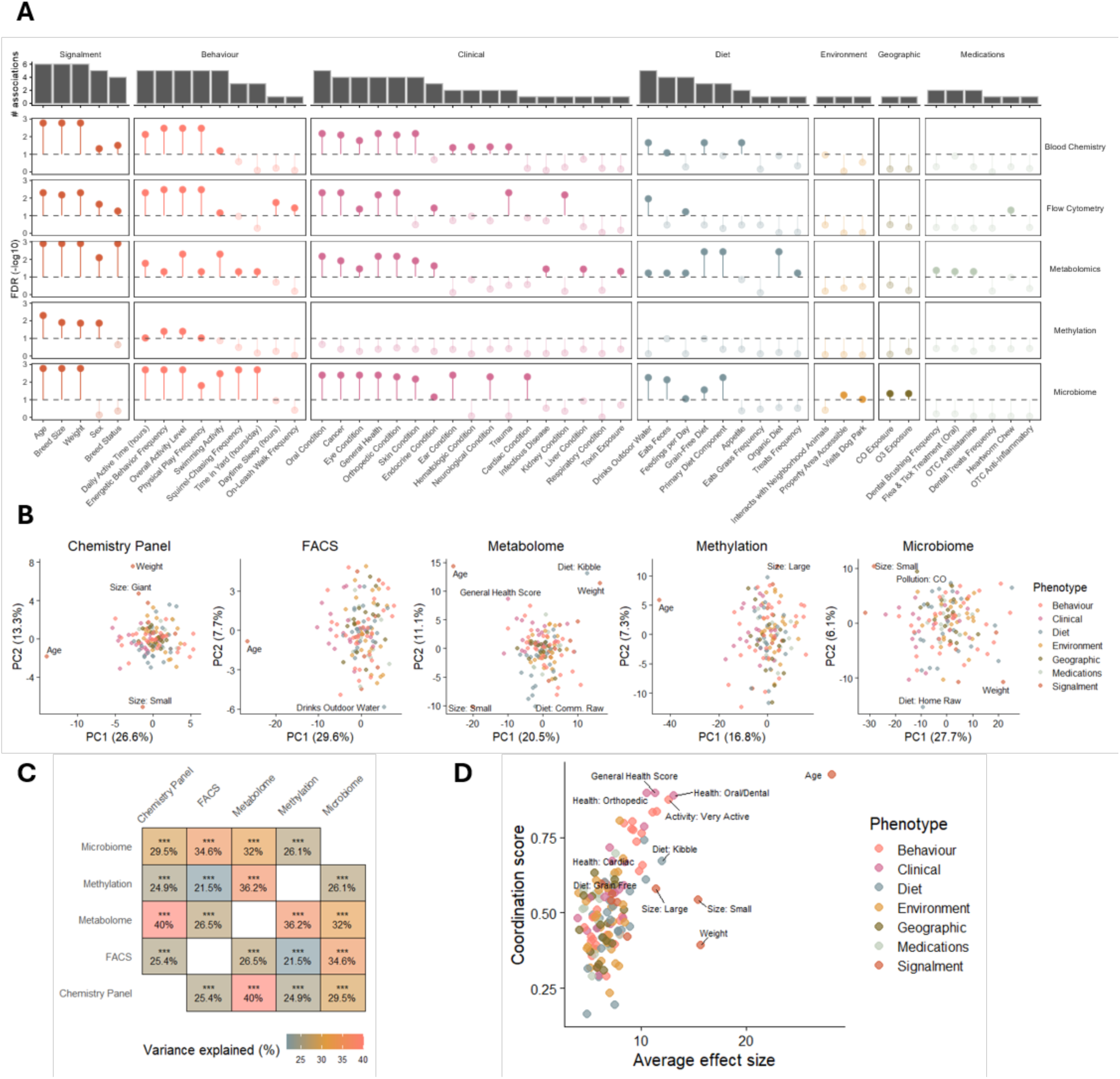
The landscape of phenotype-omic associations. **(A)** Association strength between phenotypes (categorized into seven groups: signalment, behavior, clinical, diet, environment, geographic, and medications) and five omic domains (blood chemistry, flow cytometry, metabolomics, methylation, and microbiome). Association significance was determined via PERMANOVA and is expressed as-log10(FDR). The dashed horizontal line indicates the significance threshold of FDR < 0.1. Points appearing above this line are highlighted in solid colors to denote statistically significant associations, while non-significant associations are semi-transparent. The top bar plot summarizes for each phenotype the total number of significantly associated omics. (**B**) Phenotype-omic association maps, capturing the magnitude and direction with which each omic shifts in relation to each phenotype. Each point represents a phenotypic or environmental factor, positioned according to the first two principal components. Point colors denote phenotype category and phenotypes showing extreme associations with specific omics are labeled for convenience. (**C**) Similarity between perturbation maps of different omics, quantified as variance explained by Procrustes analysis (FDR-corrected p-values). (**D**) Scatter plot comparing coordination score and average effect magnitude across phenotypes.

To further examine the magnitude and direction of such phenotype-omic effects, we constructed *phenotype-omic association maps* that aim to capture how features within each omic domain shift in relation to specific phenotypes. Specifically, for each phenotypic and environmental factor and for each omic, we computed the associations between that factor and all the features in the omic of interest. We then treated these associations as vectors (each representing the magnitude and direction with which a given omic shifts in relation to a given phenotype), and applied principal component analysis (PCA) to these vectors to assess shared trends and dominant axes of variation (Fig. 3B, Supplementary Table 7, see Methods for details). Across all omics, age consistently occupied the most extreme position along the first principal component, implicating age as the principal axis along which systemic physiological variation is structured. Perhaps unsurprisingly, clinical conditions aligned closely with this axis, reinforcing the central role of age in shaping health and disease risk. In contrast, dietary factors, for example, were largely orthogonal to age and loaded primarily along the second principal component, pointing to distinct pathways of influence. Together, these findings delineate a coherent, multi-omic axis of aging in dogs, alongside an independent, diet-driven axis of variation.

Finally, we examined the alignment between the phenotype-omic association maps generated above, to assess whether different omics shift in response to various phenotypes in the same way. This analysis revealed coordinated multi-omic alignment (Fig. 3C), with the metabolome, for example, exhibiting highly concordant association with both blood chemistry (40.0% of the variance explained; FDR < 0.001, Procrustes analysis), and methylation (36.2%). In contrast, FACS and methylation exhibited weaker concordance (21.5%), suggesting more compartmentalized associations. Moreover, quantifying the average effect size of each phenotype on the various omics, as well as the level of coordination of its effect across omics (see Methods), again highlights age as uniquely combining near-perfect coordination in its impact across the various omics with the largest effect size (Fig. 3D). Other phenotypic and environmental factors, in contrast, showed a more limited systemic association, both in terms of size and coordination. Together, these results pinpoint age as the dominant and most pervasive source of multi-omic variation in our cohort, motivating deeper analysis of age-associated signatures.

### Integrated aging clocks across multiple omics

Having established that age is a major modulator of molecular processes across multiple omic profiles, we next sought to quantify the extent to which these omics, individually or jointly, can predict chronological age and capture age-associated molecular variation. To this end, we constructed a multi-omic age clock, trained to predict a given dog’s age based on its omic profiles (see Methods). Analyses were restricted to the 533 healthy dogs (defined as owner-reported “excellent” or “very good” health) to focus on healthy aging and minimize confounding by disease and the model’s accuracy was evaluated using cross-validation. We found that the model’s predictions closely tracked true age, with a correlation of 0.817 between the mean prediction (across folds and repeats) and real age (Fig. 4A; Supplementary Table 8). Feature-importance analysis further showed that methylation, perhaps not surprisingly, contributed the most to prediction^39^, with other omics adding consistent, albeit modest contributions (Fig. 4B, Supplementary Fig. 6). Omic-specific models further confirmed the potential contribution of each omic domain, with the methylation-specific model yielding the most accurate age prediction and the microbiome-specific model the least accurate prediction (Fig. 4C).

**Figure 4.**
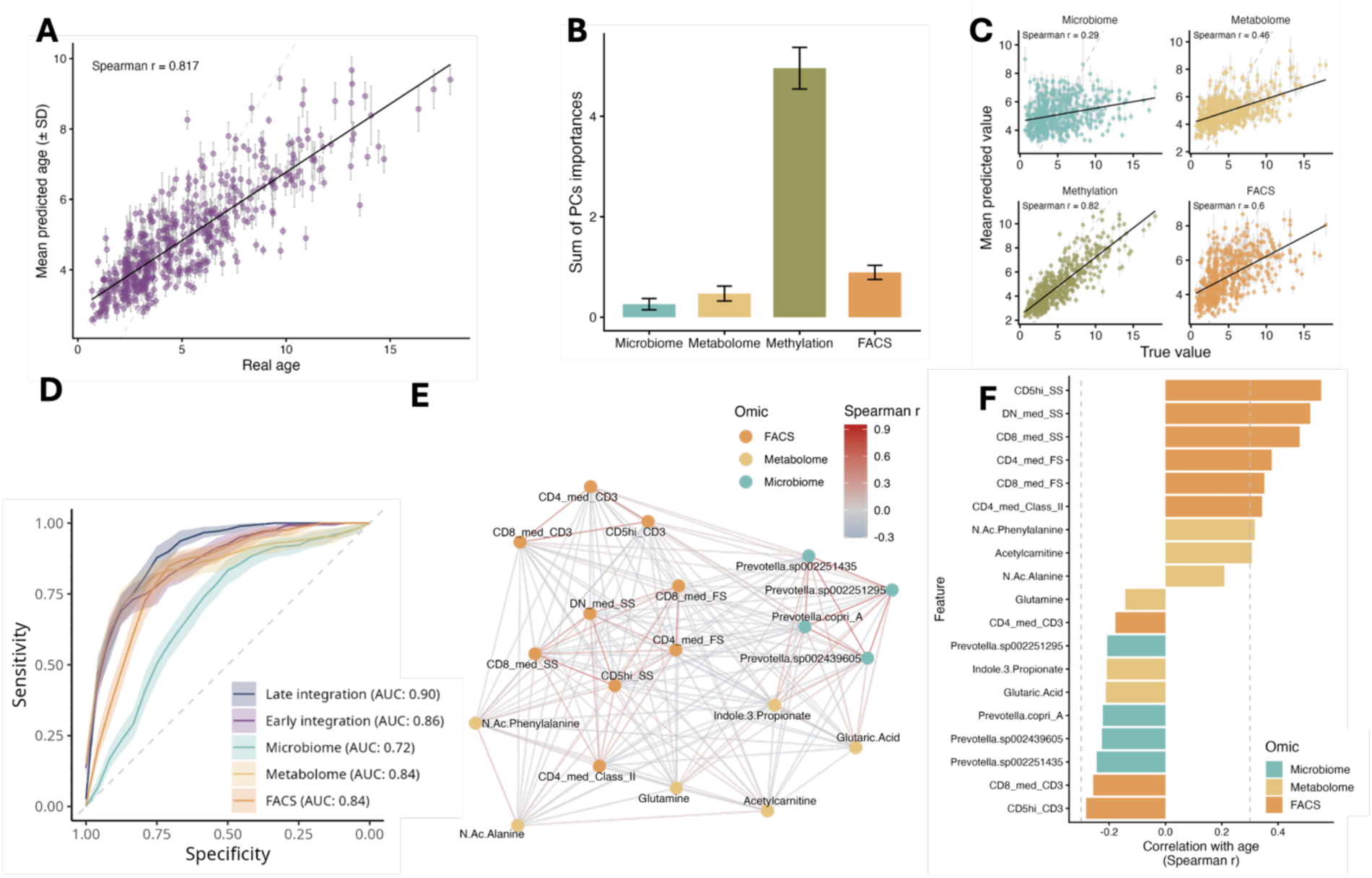
Association of multi-omic domains with age in healthy dogs. **(A**) Correlation between real and predicted age from a repeated 5-fold cross-validated model that uses the first 30 PCs from each omic. Each point is a dog’s mean predicted age, with error bars showing SD across repeats as a function of the dog’s real age. **(B)** Sum of PC importance scores for age prediction (Altmann permutation) by omic; error bars show SD across folds and repeats. **(C)** Correlation between real and predicted age, as in Panel A, but for models trained separately on each omic domain. **(D)** ROC curves for age-category prediction (young–mature vs. senior) using random forest models trained on each omic separately and two integration strategies, with mRMR feature selection (30 features) performed within each training fold. Analysis shown without methylation; the analogous plot with methylation appears in Supplementary Figure 7. **(E)** MintTea module associated with age (methylation excluded): nodes denote the various features comprising the module, colored by omic type; edges show pairwise Spearman correlations between module features, displaying only significant edges (FDR < 0.1). **(F)** Per-feature correlations with age for the module shown in panel E (bars colored by omic). Corresponding plots for analyses that include methylation are provided in Supplementary Figure 10.

We additionally evaluated the predictive capacity of the various omics when reframing the task as predicting age categories (young-mature vs. senior; see Methods) rather than continuous age. This setting facilitates a more robust analysis of predictive performance across omics with very different dimensions, distributions, and statistical characteristics. Moreover, since methylation was so highly predictive in our previous analysis, we initially excluded it from the current analysis, allowing us to assess the added value of other omic domains without methylation dominating the signal. We found that the metabolome and immunophenotyping achieved the best prediction (AUROC = 0.84), followed by the microbiome (AUROC = 0.72; Fig. 4D). Integrating omics further improved accuracy: early integration (i.e., feature concatenation) reached AUROC = 0.86, while late integration (averaging predictions from omic-specific models) performed best (AUROC = 0.90). Repeating the analysis with methylation yielded a qualitatively similar pattern (Supplementary Fig. 7).

Finally, to assess the robustness of these findings to the health-based filtration and to avoid potential survivorship bias, we repeated all age-prediction analyses using the full cohort, irrespective of health status. The obtained results were qualitatively similar throughout (Supplementary Fig. 8), including, for example, a high correlation between chronological age and age predicted by the multi-omic age clock integrating all four omic domains, now trained and evaluated using the full cohort (Spearman r = 0.831).

### A multi-omic module analysis reveals a coordinated multi-omic aging signature

While the analyses above demonstrate that multi-omic data collectively captures robust aging-related signal, it does not reveal which molecular features across the various omics might be coordinated in their association with age and might, accordingly, form a coherent multi-omic signature of aging. To address this, we used MintTea^40^, a computational framework for extracting coherent cross-omic modules whose features are both strongly correlated with each other and collectively shift with a phenotype of interest. For this analysis, we again defined age as a binary phenotype (distinguishing young-mature vs senior dogs), and ran the analysis first without methylation. In these settings, MintTea identified a single large multi-omic module, comprising 19 features in total (including 9 cell populations, 6 metabolites, and 4 microbial species), that is associated with age (Fig. 4E; Supplementary Table 9). Notably, the average cross-omic correlation between the features included in this module was significantly higher than expected under a permutation-based null model constructed from size-matched randomly sampled modules (Supplementary Fig. 9A). Consistent with its coordinated multi-omic structure, the module’s ability to distinguish dogs at different life stages (based solely on the first PC derived from the features comprising the module) was surprisingly high (AUROC = 0.88 on held-out validation data), again far exceeding that of size-matched random modules (Supplementary Fig. 9B-C). It is also important to note that not all the features in this module were strongly correlated with age on their own (Fig. 4F; Supplementary Table 10), suggesting a multivariate, synergistic, and multi-omic signature of aging that emerges at the module-level rather than from individual features. Consistent with this hypothesis, none of the individual features within the module achieved comparable discriminatory performance on their own (maximum AUROC = 0.835), suggesting that the predictive signal arises from their coordinated integration rather than from any single dominant feature.

Turning to examine the immune, metabolic, and microbial features included in this module, we further found that many of these features closely match known age-related trajectories previously observed in human cohorts. For example, several immune-cell features in the module were consistent with increased immune activation, including elevated HLA-DR expression and increased cell size and complexity in CD4+ and double negative (DN) T cells. Together, these patterns are consistent with the well-established shift from naïve T-cell dominance toward more activated and memory-skewed populations in aging^41,42^. As observed previously^32^, metabolomic signals also reflected recognized biochemical patterns, such as an age-associated rise in N-acetyl-phenylalanine, N-acetyl-alanine, and acetylcarnitine, which mirrors human reports of increased acylcarnitines and amino-acid derived metabolites with aging^43,44^. Similarly, indole-3-propionic acid (IPA), a microbiota-derived product of tryptophan metabolism, which was included in this module and negatively correlated with age, has been shown repeatedly to be lower in older adults or individuals with age-associated metabolic decline, consistent with reduced abundance of IPA-producing gut microbes^45,46^. Finally, microbial features in the module included *Prevotella copri*, as well as two additional *Prevotella* MAGs, all of which showed negative correlation with age in our cohort. Age-related shifts in *Prevotella* abundance have been reported in humans, but notably, the specific patterns varied across cohorts^47,48^. Collectively, these findings position the identified module as a coordinated immune, metabolic, and microbial aging signature, potentially capturing a universal, cross-omic axis of aging.

Repeating this analysis with the methylation data included yielded a larger module with 31 features (11 cell populations, 9 metabolites, 6 promoter-associated methylation sites, and 5 microbial species) and, as expected, higher predictive performance (AUROC = 0.95; Supplementary Fig. 10). Importantly, however, 18 of the 19 non-methylation features identified in the previous module were retained in this expanded module, indicating that the core multi-omic aging signature remains stable and that methylation contributes a largely complementary signal.

### Multi-omic trajectories of canine aging reveal size-dependent molecular temporal dynamics

Biological aging rarely proceeds at a constant pace and is often marked by discrete physiological transitions that reorganize molecular systems. In domestic dogs, adult body size is the strongest predictor of lifespan, yet it is not yet clear whether reduced longevity of large breeds reflects a uniform acceleration of aging or a more punctuated restructuring of age-dependent molecular trajectories^28,49^. Here, we set out to determine whether aging in large dogs simply represents a more compressed version of conserved aging programs or instead exhibits a reconfiguration of molecular trajectories. To this end, we implemented a feature-wise competitive modelling framework, which allowed us to determine for each molecular feature which age-size model best described its age-associated trajectory across dogs (Fig. 5A, Methods). We specifically tested whether trajectories fit a continuous linear model or one that includes a discrete structural breakpoint, and how they vary with dog size.

**Figure 5.**
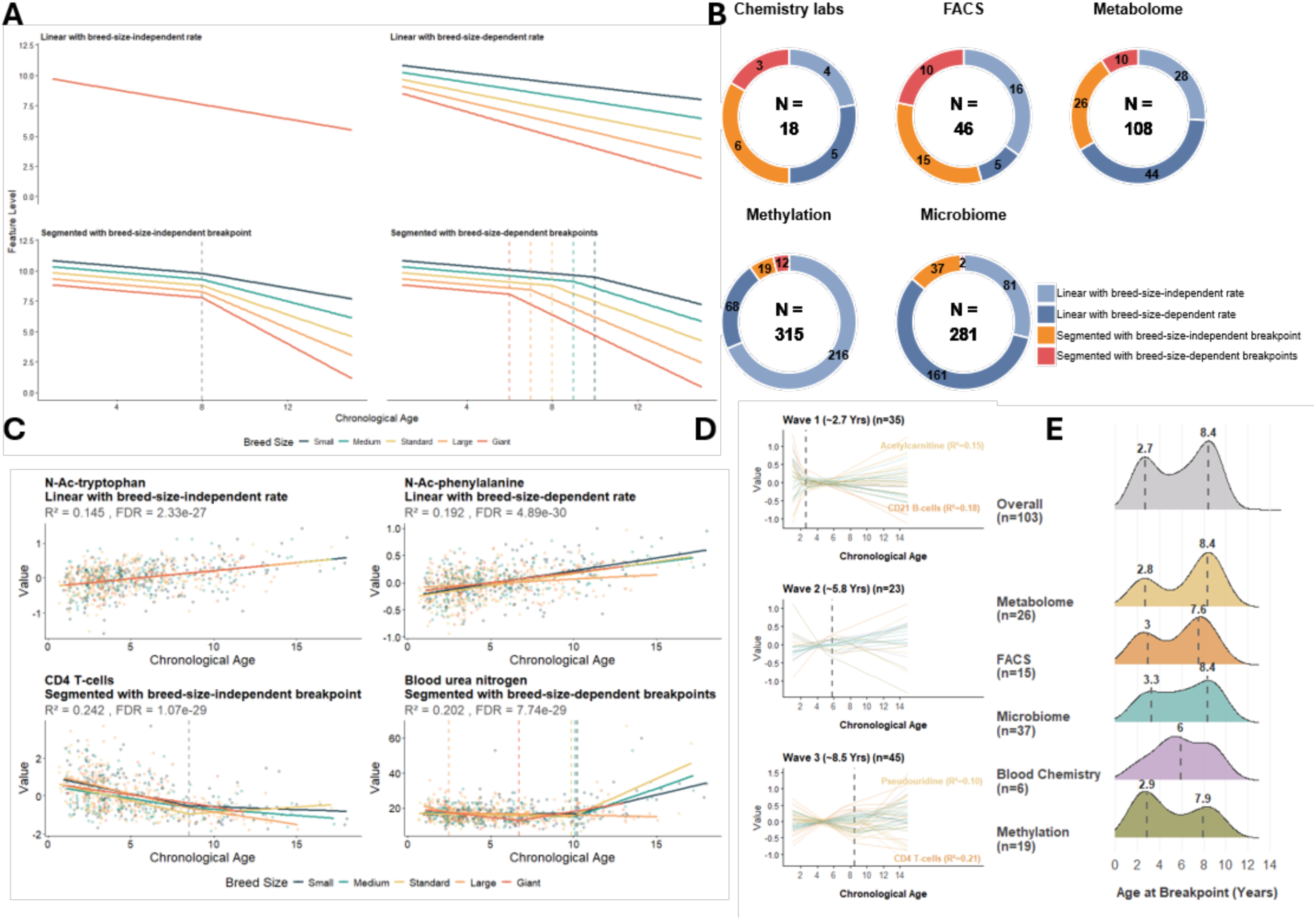
Multi-omic trajectories of canine aging reveal size-dependent molecular temporal dynamics. (**A**) Conceptual illustration of our feature-wise competitive modelling approach, used to classify molecular aging trajectories. Features are first categorized into linear versus segmented models. Linear features are further classified into two models: breed-size-independent and breed-dependent rates. Similarly, segmented features are classified into those with breed-independent and breed-dependent breakpoints. (**B**) Distribution of aging trajectory classes across the five omics domains. The number in the middle of each circle denotes the total number of significant age-associated features per domain (FDR < 0.1, F-test). (**C**) Representative examples of molecular features exhibiting distinct aging dynamics. Top: N-Ac Phenylalanine and N-Ac Tryptophan. Bottom: CD4+ T cell and blood urea nitrogen measurement. Colors indicate breed sizes. (**D**) Clustering of segmented breed-independent trajectories according to breakpoint location at approximately 2.6, 5.8 and 8.5 years of age. (**E**) Density distribution of breakpoint ages for all features exhibiting segmented breed-independent dynamics and stratification of breakpoint ages by omic domain.

Across all omic domains, 768 molecular features could be significantly explained by one of the age-size models we tested (see Methods, likelihood ratio test, FDR < 0.1, Fig. 5B, Supplementary Table 11). Of these, 628 features (∼82%) followed a linear age-associated trajectory, consistent with a continuous accumulation of molecular change across lifespan. Within this linear model class, we further distinguished between two models (Fig. 5A, top): (i) linear with breed-size-independent rate (n = 345, 45% of features), which represents an age-associated incline or decline at a consistent rate across all breed sizes, and (ii) linear with breed-size-dependent rate (n = 283, 37%), which represents breed size-dependent variation in aging rates. For example, the two N-terminally acetylated (N-Ac) amino acids phenylalanine and tryptophan, which are known to accumulate during human and canine aging^43,50^ and are associated with increased hazard ratios for cardiovascular mortality^51^, both exhibited a linear increase with age. However, while N-Ac-tryptophan appeared to accumulate at a constant rate across all breed sizes, N-Ac-phenylalanine demonstrated a breed-size-dependent trajectory, with larger dogs showing a slower accumulation with age (Fig. 5C). This difference in pace may reflect differences in microbial biosynthesis rate between the two metabolites^36,52^.

In contrast to these linear dynamics, 18% of the features (n = 140) were best described by segmented trajectories. These features exhibited a breakpoint at which the rate of change shifted, consistent with a phase-like reorganization of biological processes. Among these features, we again distinguished between two models that described the relationship between aging pace and dog size (Fig. 5A bottom): (i) segmented with breed-size-independent breakpoint (n = 103, 13% of total features), which represents a similar breakpoint age across breed sizes and (ii) segmented with breed-size dependent-breakpoints (n = 37, 5%), which exhibited a breed-specific breakpoint age. As an example, absolute counts of CD4⁺ T cells declined with age and exhibited a shared breakpoint at approximately 8.6 years of age across all breed size groups, while blood urea nitrogen levels exhibited a breed-size-dependent breakpoint, with an earlier breakpoint time in larger dogs, (6.7 vs 10.2 years for giant vs small breeds, respectively, Fig. 5C, Supplementary Table 11), reflecting earlier renal function decline in larger breeds.

We next examined the distribution of breakpoint ages across the 103 features exhibiting a segmented trajectory with breed-size-independent dynamics, to determine whether age-associated transitions occur uniformly across lifespan or cluster in discrete phases. This analysis revealed a multimodal distribution, with prominent peaks centred at approximately ∼2.6, ∼5.8 and ∼8.5 years of age (Fig. 5D) corresponding to changes in canine adulthood, mid-life and late-life. Similar clustering of age-associated molecular changes has been reported in human aging, with transitions occurring in distinct waves, one around the fourth or fifth decade of life, and another around the seventh or eighth decade^53,54^. Stratification of this distribution by omic domain further revealed pronounced differences in transition timing (Fig. 5E), with transitions tending to occur both early (showing peaks at ∼2.8-3 years) and late (around 7-8.5 years) in life. Together, these findings reveal that systemic aging in dogs is not a uniform process, but rather a complex interplay of both universally conserved linear changes and segmented biological shifts that are intricately scaled by breed size.

### Mortality prediction and kinetics of physiological decline across molecular systems

To characterize the multi-omic signatures of mortality, we leveraged the 112 death events (10.4%) observed during the follow-up period of the Precision Cohort (using the end-of-life survey; see Methods). We first evaluated how well each omic predicts survival by training omic-specific random survival forest models, using a 5-fold cross-validation framework to estimate predictive performances of each model (Methods). This analysis revealed clear differences in predictive power across the biological modalities, with mean time-dependent AUROC ranging from 0.64 for the gut microbiome to 0.7 for DNA methylation, 0.74 for the metabolome, and 0.72 for blood chemistry (Fig. 6A). Most notably, immune cell composition provided the most accurate overall predictive value (mean AUROC = 0.79), suggesting that immune-cell composition captures a strong signature of physiological decline before mortality events. Across all domains, predictive performance generally progressively decayed with longer prediction horizons, reflecting the dynamic, time-sensitive nature of these biological states (Fig. 6A). To further assess how these models stratify clinical risk, we separated individuals into high-and low-risk quartiles (highest and lowest 25%) based on their out-of-fold cross-validation predicted risk scores and performed Kaplan–Meier survival analysis (Fig. 6B). While all omic domains achieved statistically significant risk stratification, the magnitude of survival divergence varied substantially, mirroring our time-dependent accuracy assessment above. Specifically, immune cell composition and blood chemistry yielded the most profound separation in survival outcomes (log-rank P = 2.1 x 10^-10^ and 3.4 x 10^-9^, respectively), whereas separation based on the gut microbiome model was markedly weaker (albeit still statistically significant).

**Figure 6.**
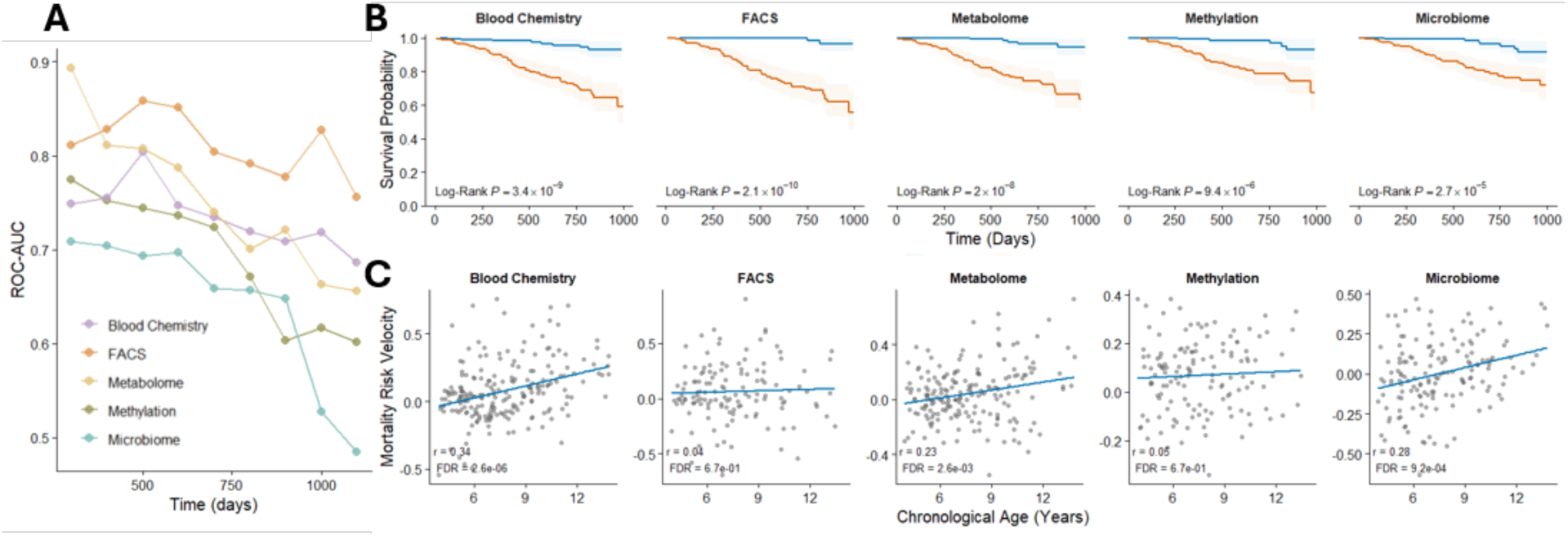
**Multi-omic integration reveals convergent signatures of aging and mortality**. (**A**) Time-dependent prediction accuracy of omic-specific survival forest models evaluated by cumulative AUROC between 200-1100 days post-sampling (10th and 90th percentiles of the observed event times). (**B**) Kaplan-Meier survival curves for dogs in our cohort stratified into high-risk (orange) and low-risk (blue) groups based on the out-of-fold predictions. The blue curve represents the top quartile and the orange curve represents the bottom quartiles of omic-specific survival predictions. Shaded regions indicate 95% confidence intervals, and P-values were determined by two-sided log-rank tests. (**C**) Dynamics of biological aging rates as a function of chronological age for each omic domain. Solid blue lines represent correlation between age and mortality risk velocity.

Interestingly, while baseline mortality risk increases exponentially with chronological age (a phenomenon classically described by the Gompertz law^55^), it remains unclear whether mortality risk associated with different omic domains increases at the same rate. To address this, we leveraged the longitudinal architecture of the Precision Cohort, which includes dogs with multi-omic measurements from two distinct timepoints, and set out to quantify the omic-specific rates of physiological decline within each subject. To this end, we derived a new metric that we termed *Mortality Risk Velocity (*ℳ), defined as the within-subject change in restricted mean survival time (RMST, which represents the average time an individual is expected to live during the follow-up window as predicted by the omic of interest) between two time points (see Methods). In this metric, a positive value denotes an increase in mortality risk with time, with higher values indicating a faster increase in risk. Evaluating mortality risk velocity against the dog age at the first omic measurement, and focusing on mature adults or senior dogs, we found an increase in mortality risk velocity as dogs became older only based on the blood chemistry, microbiome, and metabolome profiles (Fig. 6C; FDR < 0.1, Pearson correlation), suggesting that these omic domains may serve as leading indicators of systemic physiological decline. In contrast, methylation and FACS did not exhibit a correlation between mortality risk velocity and age. Together, these findings demonstrate that physiological aging is not a uniform systemic deterioration, but rather a composite of unique molecular trajectories, with specific subsystems driving acceleration of mortality risk.

## Discussion

In this study, we leveraged a longitudinal, deeply phenotyped, multi-omic cohort of 965 companion dogs to gain a systems-level view of canine aging. We found a strong, prevalent, yet structured cross-omic concordance, with particularly tight coupling between the microbiome and the metabolome and between epigenetic and immune profiles. Systematic mapping of phenotype-omic associations highlighted age as the dominant and most coordinated axis of biological variation, overshadowing lifestyle and environmental factors, which exerted more localized, omic-specific effects. Integrative modeling further demonstrated that chronological age is most strongly encoded in the epigenome, with the immune, metabolic, and microbial domains providing complementary and partially independent aging signals that jointly improve age prediction. We also found a coherent cross-omic aging module that integrates immune, metabolic, and microbial features into a unified aging signature. Feature-wise trajectory analyses further showed that aging is manifested through both continuous and segmented molecular dynamics, with breed size modulating baselines, rates, and, more rarely, the timing of age-associated transitions. These analyses further revealed discrete waves of molecular reorganization emerging in midlife and later adulthood. Finally, longitudinal survival analysis suggested that aging is a heterogeneous process and demonstrated that distinct molecular domains encode different aspects of vulnerability and robustness during aging. Importantly, although the epigenome reflects chronological age, metabolic and immunophenotypic profiles serve as superior predictors of mortality, highlighting how different omic domains capture distinct biological processes.

Notably, several of the patterns observed above mirror those reported in human cohorts, pointing to conserved organizing principles of mammalian aging. For example, the cross-omic coupling reported above recapitulates well-established relationships in humans, where microbial metabolic activity strongly influences circulating metabolite profiles^56^ and epigenetic variation in blood is tightly coupled to immune cell composition and age-related immunological remodeling^57,58^. Consistent with this conservation, multiple metabolites that are robustly predictable from the microbiome in dogs, including indole-3-propionate, 3-indoxyl sulfate, phenylacetylglutamine, and related amino acid-derived compounds, exhibit similar age-associated behavior and disease relevance in large human cohorts^59–61^. Moreover, the microbiome-based predictability of each metabolite in dogs (out-of-sample predicted–measured correlation) was correlated with the proportion of variance explained by the microbiome in human serum^37^, implicating shared host-microbe metabolic pathways across species. At the integrative level, the predominance of DNA methylation in age prediction, alongside additional and partially orthogonal contributions from immune and metabolic domains, parallels findings from human multi-omic aging studies, where epigenetic clocks capture the strongest chronological signal while other omics encode complementary dimensions of biological aging^62,63^. Similarly, aging in dogs, as in humans, appears not to follow a temporally uniform pace: the clustering of molecular transitions around midlife and later adulthood aligns with reports of discrete waves of molecular and physiological reorganization in humans, rather than gradual linear change^53,64^. Extending this concordance to survival-related analysis, we again found many parallel trends, including, for example, age-associated declines in Holdemanella species, and the marked impact of blood-based domains (chemistry and immune state) on survival (although human evidence to date has predominantly come from single-omic or partially integrated studies^65,66^). Together, these parallels support the use of companion dogs as a translational model that captures both the structure and dynamics of human aging in a real-world physiological context.

Clearly, several sources of bias and methodological constraints should also be considered when interpreting our findings. As in other large prospective cohorts, survivor bias is possible^67^: dogs who remain enrolled at older ages may represent a particularly resilient subset, potentially leading to an underestimation of aging-related deterioration. The cohort is also subject to self-selection, as participation depends on motivated owners, and dogs with significant health or behavioral issues may be underrepresented. Combined with the fact that all participants were recruited in the United States, these factors suggest that some of the observed patterns may reflect healthier and more environmentally homogeneous aging trajectories than would be found in the global dog population. Additionally, owner-reported health status, while informative^68^, may introduce additional noise into phenotypic classifications. Previous analyses of the Dog Aging Project have also identified owner age as a significant predictor of several owner-reported measures, including perceived health and physical activity, suggesting that owner characteristics may contribute to variation in survey-derived phenotypes^69^.

From a methodological standpoint, multi-omic integration introduces several challenges. Different data types vary markedly in dimensionality, sparsity, and noise characteristics, which can influence the stability of learned cross-omic relationships^70,71^. For example, the various omics analyzed in this study differ markedly in feature dimensionality; methylation includes tens of thousands of sites, whereas FACS has relatively few features. Such imbalances can cause high-dimensional omic domains to dominate shared structure, potentially obscuring signals from lower-dimensional omics. We mitigated this in multiple analyses (e.g., using PCA-based dimension reduction, factor modeling, or per-omic normalization), yet some residual influence is unavoidable. Second, batch effects and technical variability differ across platforms and are difficult to fully harmonize in a dataset of this scale.

Ultimately, however, this work establishes a longitudinal, multi-omic resource that enables aging to be studied as a dynamic, system-level process in a large population of companion animals at a scale and resolution that has not previously been introduced. By repeatedly profiling microbial, metabolic, epigenetic, immune, and clinical domains in dogs living in human environments, the Precision Cohort allows biological trajectories to be examined directly over time rather than inferred from cross-sectional contrasts. This enables integrated analyses that connect molecular change to physiology, environment, and health status, while preserving the complexity of real-world aging. More broadly, this framework supports a conceptual shift from viewing aging as a single linear process to understanding it as a multidimensional phenomenon emerging from interactions among biological systems and shaped by genetics, environment, and life history. By making these data and analytical approaches publicly available, this work provides a foundation for identifying molecular signatures that precede overt disease or functional decline, informing the design and evaluation of interventions targeting age-associated processes, and strengthening the translational bridge between companion animal models and human geroscience while explicitly accommodating the heterogeneity and contingency inherent to aging biology.

## Methods

### DAP Precision cohort description

The Dog Aging Project (DAP) is a large, nationwide longitudinal study designed to investigate the biological and environmental factors that shape aging in companion dogs. Within this initiative, the Precision Cohort represents a deeply phenotyped subset of dogs for whom extensive multi-omic measurements were generated^29^. This cohort includes 976 dogs with stool metagenomic profiles, blood-based metabolomics, whole-genome sequencing, methylation data, and detailed immunophenotyping. Routine clinical laboratory measurements, such as complete blood counts, chemistry panels, and urinalysis, were also obtained, and owners provided comprehensive information on health, lifestyle, and environment through structured surveys administered at enrollment and annually thereafter. In the present study, we primarily analyzed the Precision dataset, retaining 965 dogs after quality control. This dataset includes baseline samples from all enrolled dogs as well as second-year samples for a subset (662 dogs), enabling evaluation of both cross-sectional and limited longitudinal structure in the data.

### Data generation and preprocessing

#### Stool microbiome

Stool specimen collection, DNA extraction, library preparation, and metagenomic shotgun sequencing were performed as previously described^29,31^. Briefly, stool samples were collected from companion dogs, processed using standardized extraction protocols, and sequenced to generate high-depth metagenomic profiles.

Microbiome data preprocessing was performed following a framework similar to that described previously^31^. Briefly reads were quality-filtered, host reads removed, samples depth-normalized, and taxonomic and functional profiles inferred, with rare taxa excluded. Relative-abundance profiles were used for all downstream microbiome analyses unless otherwise noted.

#### Serum metabolome

Metabolomic profiling was performed as previously described^29,32^. Briefly, aqueous metabolites were extracted from plasma samples and quantified using targeted liquid chromatography–tandem mass spectrometry (LC–MS/MS). This assay yielded peak-area measurements for 133 metabolites.

Preprocessing of metabolomic data was conducted following a workflow similar to that described previously^32^, with details provided here. Raw peak-area values were log-transformed and mean-centered. To account for technical variation, metabolite abundances were adjusted for extraction batch, LC–MS run order, sample hemolysis, sample lipemia, and sample arrival temperature. Missing values were imputed using a custom elastic-net–based procedure, as implemented in a previously described pipeline^32^.

#### DNA methylation

PBMC DNA methylation data were generated and processed as previously described^29,33,34^. Promoter regions were subsequently defined as the 2 kb upstream of transcription start sites (TSS) based on the CanFam4 reference genome (https://www-ncbi-nlm-nih-gov.ezproxy1.lib.asu.edu/datasets/genome/GCF_011100685.1/). Consistent with standard practice for autosomal methylation analyses, loci on the X and Y chromosomes were excluded to avoid sex-specific methylation effects.

#### Flow cytometry

Flow cytometry (FACS) data were obtained as described before^29^. We first assessed feature-and sample-level missingness, excluding dogs with more than 20 missing FACS markers and removing three markers with more than 10 missing values. The remaining eight missing values across five markers were imputed using k-nearest neighbors. To correct for temporal technical drift, each marker was modeled as a smooth function of runDay using spline regression (R package ‘splines’ version 4.2.2), and the residuals were used as drift-adjusted values, following principles established for drift correction in cytometry and other omic assays^72,73^. Outliers were then identified based on multivariate distance in PCA space: principal components were computed from all drift-adjusted markers, Euclidean distances from the PCA centroid were calculated using the first five PCs, and samples exceeding a threshold of mean + 3 SD were removed. Finally, all markers were standardized to zero mean and unit variance to ensure consistent scaling across features in downstream analyses.

### Global association between omics

To estimate the global associations between omic domains, we first quantified pairwise cross-omic concordance using sparse canonical correlation analysis (sCCA). Briefly, for each pair of omic domains we restricted the analysis to dogs profiled in both domains and z-score standardized each feature using parameters estimated from the training samples and applied to the corresponding held-out samples. We then fit sCCA models using the PMA package (version 1.2.4), using 25 permutations within each training set to select the L1 penalties by comparing the observed canonical correlation with correlations obtained after permuting the correspondence between the two omic datasets, and evaluating the resulting models using a 5-fold cross-validation scheme (using the first canonical component). For each fold, we computed the Pearson correlation and corresponding P-value between the first pair of canonical variates in the held-out samples and summarized cross-validated concordance as the mean correlation across folds. The five-fold-specific P-values were combined using Fisher’s method and adjusted for multiple testing using the Benjamini–Hochberg false discovery rate (FDR). For visualization, the squared mean cross-validated canonical correlation (r²) was plotted. As an additional validation of these cross-omic relationships at the global level, we computed between-dog distance matrices for each data type. Distance matrices were constructed as follows: microbiome profiles were compared using Bray-Curtis dissimilarity applied to species-level relative abundances; genotype distances were based on the genetic relationship matrix computed with PLINK; and Euclidean distances were used for all other omic domains. Pairwise covariation between distance matrices was quantified using the Procrustes test, implemented with the ‘protest’ function from the vegan package (version 2.6-4) with 999 permutations. P-values were adjusted for multiple testing using the Benjamini-Hochberg false discovery rate (FDR).

### Metabolites and immune cells prediction

We trained random forest models in a 5-fold cross-validation scheme to predict metabolite abundances from microbiome features and immune-cell feature abundances from methylation features, using the ranger package (version 0.16.0). Methylation analyses were restricted to highly variable CpG sites (variance > 0.01 across samples). For each predicted feature, prediction accuracy was evaluated in held-out test folds using Spearman correlation between observed and predicted values. Correlation coefficients and corresponding p-values were obtained for each fold and then summarized across folds by computing the mean and standard deviation of the correlations. To combine evidence across folds, p-values were aggregated using Fisher’s method, and multiple testing correction was done using the Benjamini-Hochberg procedure. Features were considered significantly predictable if the mean correlation between their real and predicted values was > 0.3 with q-value < 0.1. Here and throughout the text, FDR <α is used to indicate tests whose corrected P values (q values) passed the specified threshold.

To compare microbiome-metabolite associations in dogs with analogous trends in humans, we used serum metabolomics data from Dekkers et al.^37^, generated in 8,583 participants of the SCAPIS cohort. For each metabolite, we extracted the variance explained (R²) by gut microbial composition from Supplementary Data 5, based on nested cross-validated ridge regression models. We compared these human association scores with our cohort’s mean cross-validated predictability (correlation between observed and microbiome-predicted metabolite levels).

To identify which microbial features contributed most to metabolite prediction, we further applied Altmann’s permutation feature importance procedure^74^, which repeatedly permutes feature values to estimate their contribution to model accuracy in a statistically rigorous manner.

### MOFA+ integration

We used MOFA+ (version 1.8.0)^38^ to derive a shared latent representation across four omic domains: centered log-ratio (CLR)-transformed species-level microbiome relative abundances, targeted metabolomics, DNA methylation, and FACS cell-population frequencies. Samples from both sampling time points were included in the analysis. For methylation, low-variance loci were removed, retaining sites with variance > 0.05 across samples, to avoid feature imbalance between the omics. During preparation we enabled per-view scaling (scale_views = TRUE) to balance differences in dimensionality and variance across views, and otherwise used default model and training options. The trained model yielded latent factors (sample scores) and feature loadings per view for downstream analyses.

### Feature-wise stability

For each omic, we computed the Spearman correlation (r) between Precision 1 and Precision 2 for every measured feature across dogs and summarized the distribution as mean ± s.d. Differences between feature-wise correlation distributions across omics were tested with paired Wilcoxon signed-rank tests; p-values were FDR-adjusted.

### Global cross-time concordance

For each omic and in the MOFA+ integrated latent factors, we constructed sample-by-sample distance matrices at P1 and P2 and ran Procrustes tests as described above in the “Global association between omics” section.

### Self-distance versus random baseline

For each dog, the self-distance was defined as the distance between its P1 and P2 profiles (Bray-Curtis for the microbiome; Euclidean for other omics). As a baseline, we calculated the mean distance between that dog’s P1 profile and the P2 profiles of all other dogs. Self-and other-dog distances were compared separately for each omic using linear mixed-effects models, with distance type (self vs. other), breed size, sex/sterilization class, and life-stage category as fixed effects and a random intercept for dog to account for the two distance measurements contributed by each dog. Dogs with missing distance or covariate values were excluded. The same analysis was applied to distances in the MOFA+ latent space, and p-values for the self-versus-other contrast were FDR-corrected across the microbiome, metabolome, DNA methylation, FACS, and MOFA+ latent-space analyses.

### Demographic contrasts

Within each omic, and in the MOFA+ integrated latent factors, we ranked P1 to P2 distances across dogs (rank 1 = smallest change). A cross-omic mean rank was then computed by averaging per-dog ranks across omics to assess overall longitudinal change. Ranks were compared across categories (e.g., breed size, sex/sterilization, life stage, health status) using Wilcoxon tests for two-level contrasts and pairwise Wilcoxon tests for multi-level factors, with FDR correction.

### Global associations between exposures and omics

To quantify the overall effect of each phenotype on each omic domain from Precision 1, we applied permutational multivariate analysis of variance (PERMANOVA), using the vegan R package (version 2.6-4)^75^ and significance was assessed using permutations followed by FDR correction.

### Phenotype-omic association maps

To characterize the directionality of perturbations, we constructed phenotype perturbation maps. For each categorical exposure, we applied one-hot-encoding and calculated feature-wise associations within each omic using linear models. This yielded a vector of effect sizes representing the perturbation profile of that exposure within an omic. Vectors were then subjected to principal component analysis (PCA) to obtain an ordination of exposures. PCA loadings were retained to identify the omic features most strongly driving each perturbation. To find whether the perturbation maps of different omics were aligned we used Procrustes analysis. For each pair of omic domains, the PCA ordinations of phenotypes were compared, and similarity was quantified by the Procrustes variance explained.

### Multi-omic coordination analysis

To evaluate systemic biological responses to various phenotypes, we implemented a multi-omic integration framework driven by two primary metrics. The first, average effect size, was designed to quantify the overall strength and systemic effect size of the phenotype across the omic domains. The second, coordination score, was designed to evaluate directional synchronization, specifically, whether different omic domains shift in a consistent direction in association with a given phenotype. For each omic domain, we extracted the first 10 principal components (PCs) and represented each phenotype as a vector in the resulting 10-dimensional PC space. This yielded one PC-based representation per omic domain. We quantified the effect size within each omic domain as the L2 norm of the corresponding PC vector and defined the average effect size as the geometric mean of these L2 norms across all omic domains. Lastly, to quantify the coordination score for each phenotype, we computed the mean cosine similarity between each individual omic PC vector and the centroid vector (the mean vector of all omic domains). A score of 1 means perfect directional alignment, 0 means orthogonality, and negative values indicate opposing directions.

### Multi-omic age prediction using principal component representations

We constructed a multi-omic age clock using a repeated 5-fold cross-validation scheme with 5 repeats. In every fold, we first harmonized samples across all four omic tables and the metadata, retaining only dogs present in all datasets and with complete feature data. We then fit PCA separately within each omic domain on the training samples only (prcomp from stats version 4.2.2, with centering and scaling). For each omic, we retained up to 30 principal components (PCs), capped by the number of samples and features, and projected the held-out test samples onto these training-derived PCs. The resulting per-omic PC scores were concatenated and combined with covariates (breed size, sex, and weight) to form a joint feature set, which was used to train a random-forest regression model (ranger version 0.16.0) to predict chronological age. Performance in each test fold was quantified by the Spearman correlation between predicted and observed age, and fold-level p-values were combined across folds using Fisher’s method. When computing feature importance, we applied permutation-based (Altmann) importance at the PC level within each fold (ranger::importance_pvalues(method = “altmann”)); we then summarized both the sum of PC importances per omic and the per-PC contributions as mean ± SD across folds and repeats. To assess the age-predictive capacity of each omic domain independently, we repeated the same cross-validation and modeling procedure separately for each omic, using only the PCs derived from that single omic, together with the same covariates (breed size, sex, and weight), as input features.

### Age class prediction from individual and integrated omics

To evaluate the predictive power of each omic domain for age, we used data from healthy dogs only and performed binary classification of age categories (young-mature and senior). For each omic dataset, we applied minimum redundancy maximum relevance (mRMR) feature selection within each training fold to retain the top 30 features, and repeated the procedure 10 times (mRMR package, version 2.1.2.1). Random forest classifiers were trained using 5-fold cross-validation, with breed size, weight, and sex included as covariates in all models (caret). For the microbiome, sequencing depth and duplication rate of the sequencing were also included as additional features in the microbiome table, and consequently were incorporated into the early integration analysis. Two multi-omic integration strategies were evaluated: (i) early integration, in which selected features from all omics together with the covariates were concatenated into a single model, and (ii) late integration, in which predicted probabilities from the omic-specific models were averaged to generate a final prediction. Analyses were repeated excluding epigenome data to assess the impact of methylation on integrated model performance.

### Identify age-associated multi-omic modules

To identify age-associated multi-omic modules, we used MintTea (version 1.0)^40^, an integration framework for multi-omic microbiome data, which combines an extension of sparse generalized canonical correlation analysis (sGCCA), consensus analysis, and an evaluation protocol to identify coherent multi-omic modules associated with phenotype. Each module consists of features from different omic domains that are both cross-correlated and collectively associated with the phenotype. This approach enables the detection of biologically interpretable, disease-associated modules that capture interactions across omic domains. We consider age as two age categories: senior vs. young-mature. We concatenated all omic features tables and used MintTea with the following parameters: param_diablo_keepX = 10 (default), param_sgcca_design = 0.7, param_edge_thresholds = 0.8 (default) and n_evaluation_folds = 5 (default). We used MintTea function plot_module_stats to evaluate the AUROC of the module’s first PC in age prediction and the average cross-omic correlation between features.

### Modeling of age-and size-dependent molecular trajectories

To characterize the relationship between chronological age, breed size according to the American Kennel Club (AKC) classification, and molecular change across multiple omic domains, we evaluated four distinct mathematical models of aging for each feature. For each feature, we tested:

(1) Linear with breed-size-independent rate model (*Y* ∼*Age*), representing a universal, constant rate of change.
(2) Linear with breed-size-dependent rate model (*Y* ∼*Age* × *Breed*_*size*), representing a constant rate of change that diverges between sizes.
(3) Segmented with breed-size-independent breakpoint model, hypothesizing a conserved biological shift in aging trajectory occurring at the same chronological age across all breeds, but with size-specific rates of decline thereafter. This model is defined as:

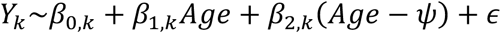

Where the breakpoint, *ψ*, is a single, shared value across all breed size categories (*k*).

(4) Segmented with breed-size-dependent breakpoints model, hypothesizing an allometric scaling of aging, where both the timing of the shift and the subsequent rate of decline are uniquely determined by breed size. This model allows the breakpoint, *ψ_k_*, to be explicitly estimated for each specific breed size category (*k*), defined as:

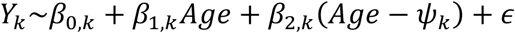

In both segmented models, the term *β*_2,*k*_(*Age* − *ψ*) or *β*_2,*k*_(*Age* − *ψ_k_*) represents the hinge function, evaluating to *Age* − *ψ* when a dog’s age exceeds the respective breakpoint, and 0 otherwise.

Segmented regression was performed using the “segmented” R package (v.2.2.0). To determine the most parsimonious fit, we utilized a hierarchical likelihood-ratio test framework. The presence of a significant non-linear breakpoint was first assessed using the Davies test (p<0.05)^76^. If a breakpoint was supported, models (3) and (4) were compared via an F-test to determine if size-dependent breakpoints significantly improved model fit over a shared breakpoint. If the Davies test failed to reject linearity, models (1) and (2) were compared similarly. To ensure the selected model represented a biologically meaningful relationship rather than stochastic noise, we applied a global F-test comparing the winning model to an intercept-only null model, retaining only features that passed an FDR<0.1 threshold. For this analysis we used CLR-transformed microbiome profiles.

### Survival-Informed Machine Learning Framework

Mortality data are based on owner-reported end-of-life surveys and include deaths that occurred from May 2019 to December 2024. For prognostic evaluation, omic-specific survival forest models were trained to predict mortality. Models were trained on baseline profiles to predict survival outcomes (follow-up time and mortality status). To ensure robust estimation and prevent overfitting, we employed 5-fold cross-validation, stratified by mortality status. Model performance was quantified using time-dependent cumulative area under the receiver operating characteristic curves (AUROC). Metrics were computed at regular intervals from 200 to 1,100 days post-sampling, capturing both short-and long-term predictive accuracy. Models were evaluated using fivefold cross-validation, and performance metrics were averaged across folds.

### Calculation of the mortality risk velocity

We used Restricted Mean Survival Time (RMST) as a quantitative proxy for biological health and physiological resilience. Unlike hazard ratios, which describe relative risk, RMST provides an absolute measure of expected remaining life up to a defined time horizon. To capture the dynamics of physiological decline, we defined a new metric termed *mortality risk velocity*, representing the change in predicted life expectancy between consecutive sampling points (and normalized by the chronological time gap between these two points). The metric is based on the change in RMST between two consecutive omics measurements (i.e. P1 and P2). At each timepoint, a survival model estimates the RMST up to a 1,300-day horizon. Formally, the mortality risk velocity, ℳ, for a subject between two timepoints was defined as:

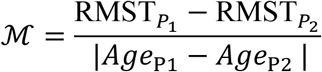

Where RMST_P1_ and RMST_P2_ are the RMST at measurements P1 and P2, and Age_P1_ and Age_P2_ are the subject’s ages at these timepoints. To characterize how mortality risk velocity changes across lifespan, we focused on mature adult or senior subjects (since these age groups have higher mortality risk) and calculated the Pearson correlation between mortality risk velocity and age at the first omic measurement (i.e., age at P1).

### Human and animal subjects research approval

The University of Washington IRB deemed that recruitment of dog owners for the Dog Aging Project, and the administration and content of the DAP Health and Life Experience Survey (HLES), are human subjects research that qualifies for Category 2 exempt status (IRB ID no. 5988, effective 10/30/2018). All study-related procedures involving privately owned dogs were approved by the Texas A&M University IACUC, under AUP 2021-0316 CAM (effective 12/14/2021).

## Data and code availability

The raw metagenomic sequencing data generated in this study have been deposited in the NCBI Sequence Read Archive (SRA) under BioProject accession PRJNA1377461. The raw metabolomics data and the code used for their normalization are available at https://github.com/ben6uw/Dog_Aging_Project_metabolome_2026/tree/main. The epigenomic sequencing data are available through the NCBI Gene Expression Omnibus (GEO) under accession GSE306794. Processed metagenomic, metabolomic, epigenomic, and FACS data, together with all scripts used for statistical analyses and figure generation, are available at https://github.com/dogagingproject/Dog-Aging-Project-Publication-Resources/tree/main/2026_bamberger_multi_omic_longitudinal_profiling_of_companion_dog. Dog Aging Project data are available through the Terra platform (https://app.terra.bio/); instructions for accessing the open data are provided on the Dog Aging Project website (https://dogagingproject.org/data-access).

## Acknowledgements

The authors thank Dog Aging Project participants, their dogs, and community veterinarians, as well as all Dog Aging Project Team members (2025) [https://dogagingproject.org/team-acknowledgements] for their important contributions. We also thank all current and past members of the Borenstein lab for their advice and help, and the IT team of TAU School of Computer Science and AI for technical support during the computational work.

## Funding

This research is based on publicly available data collected by the Dog Aging Project, under U19 grant AG057377 (PI: Daniel Promislow) from the National Institute on Aging, a part of the National Institutes of Health, and by additional grants and private donations, including generous support from the Glenn Foundation for Medical Research, the Tiny Foundation Fund at Myriad Canada, the WoodNext Foundation, and the Dog Aging Institute. These data are housed on the Terra platform at the Broad Institute of MIT and Harvard. This work was further supported by the Israel Science Foundation grant 2266/25 to E.B. T.B. is supported in part by a fellowship from the Edmond J. Safra Center for Bioinformatics at Tel-Aviv University. Y.M.A. is supported in part by fellowships from the Foulkes Foundation and the Dog Aging Institute. The content is solely the responsibility of the authors and does not necessarily represent the official views of the National Institutes of Health.

## Declaration of interests

N.S.-M. is an inventor on a pending US patent application related to this work (Canine Epigenetic Clock and Related Methods; US Patent Application no. 63/891,072; filed 30 September 2025). D.P. is founder and treasurer of the Dog Aging Institute and serves on the Scientific Advisory Boards of WNDRHlth and the Morris Animal Foundation. The other authors declare no competing interests.

## The Dog Aging Project Consortium

Joshua M. Akey^a^; Rozalyn M. Anderson^b^; Anne Avery^c^; Elhanan Borenstein^d^; Marta G. Castelhano^e^; Frances L. Chen^f^, Weihsueh A. Chiu^g^; Amanda E. Coleman^h^; Laura Corlin^i^; Kate E. Creevy^j^; Matthew D. Dunbar^k^; Virginia R. Fajt^l^; Jessica M. Hoffman^m^; Matt Kaeberlein^n^; Elinor K. Karlsson°; Kathleen F. Kerr^p^; Evan L. MacLean^q^; Stephanie McGrath^r^; Natasha J. Olby^s^; Daniel E. L. Promislow^t^; May J. Reed^u^; Audrey Ruple^v^; Stephen M. Schwartz^w^; Sandi Shrager^x^; Noah Snyder-Mackler^y^

^a^Lewis-Sigler Institute for Integrative Genomics, Princeton University, Princeton, NJ, USA; ^b^Department of Medicine, University of Wisconsin–Madison, Madison, WI, USA; ^c^Department of Microbiology, Immunology and Pathology, College of Veterinary Medicine and Biomedical Sciences, Colorado State University, Fort Collins, CO, USA; ^d^Department of Clinical Microbiology and Immunology, Gray Faculty of Medical and Health Sciences, Tel Aviv University, Tel Aviv, Israel; ^e^Cornell Veterinary Biobank, College of Veterinary Medicine, Cornell University, Ithaca, NY, USA; ^f^Genomics and Computational Biology, UMass Chan Medical School, Worcester, MA; ^f^Department of Small Animal Medicine and Surgery, College of Veterinary Medicine, University of Georgia, Athens, GA, USA; ^g^Department of Veterinary Physiology and Pharmacology and Interdisciplinary Faculty of Toxicology, College of Veterinary Medicine & Biomedical Sciences, Texas A&M University, College Station, TX 77843; ^i^Tufts University, Public Health and Community Medicine, School of Medicine; ^j^Department of Small Animal Clinical Sciences, Texas A&M University School of Veterinary Medicine & Biomedical Sciences, College Station, TX, USA; ^k^Center for Studies in Demography and Ecology, University of Washington, Seattle, WA, USA; ^l^Department of Veterinary Physiology and Pharmacology, Texas A&M University School of Veterinary Medicine & Biomedical Sciences, College Station, TX, USA; ^m^Department of Biological Sciences, Augusta University, Augusta, GA, USA; ^n^Department of Laboratory Medicine and Pathology, University of Washington School of Medicine, Seattle, WA, USA; °Bioinformatics and Integrative Biology, University of Massachusetts Chan Medical School, Worcester, MA, USA; ^p^Department of Biostatistics, University of Washington, Seattle, WA, USA;^q^College of Veterinary Medicine, University of Arizona, Tucson, AZ, USA; ^r^Department of Clinical Sciences, College of Veterinary Medicine and Biomedical Sciences, Colorado State University, Fort Collins, CO, USA; ^s^Department of Clinical Sciences, College of Veterinary Medicine, North Carolina State University, Raleigh, NC, USA; ^t^Jean Mayer USDA Human Nutrition Research Center on Aging at Tufts University, Boston, MA, USA; ^u^Department of Medicine, Division of Gerontology and Geriatric Medicine, University of Washington School of Medicine, Seattle, WA, USA; ^v^Department of Population Health Sciences, Virginia-Maryland College of Veterinary Medicine, Virginia Tech, Blacksburg, VA, USA; ^w^Epidemiology Program, Fred Hutchinson Cancer Center, Seattle, WA, USA; ^x^Collaborative Health Studies Coordinating Center, Department of Biostatistics, University of Washington, Seattle, WA, USA; ^y^School of Life Sciences, Arizona State University, Tempe, AZ, USA

## Supplemental information

Document S1. Supplemental text and Figures S1-S10.

Data S1. Tables S1-S11.

## Notes

https://github.com/dogagingproject/Dog-Aging-Project-Publication-Resources/tree/main/2026_bamberger_multi_omic_longitudinal_profiling_of_companion_dog

## References

1. Hou, Y. et al. Ageing as a risk factor for neurodegenerative disease. Nat. Rev. Neurol. 15, 565–581 (2019).

2. Kaeberlein, M., Rabinovitch, P. S. & Martin, G. M. Healthy aging: The ultimate preventative medicine. Science 350, 1191–1193 (2015).

3. Abdellatif, M., Rainer, P. P., Sedej, S. & Kroemer, G. Hallmarks of cardiovascular ageing. Nat. Rev. Cardiol. 20, 754–777 (2023).

4. López-Otín, C., Blasco, M. A., Partridge, L., Serrano, M. & Kroemer, G. Hallmarks of aging: An expanding universe. Cell 186, 243–278 (2023).

5. Manolio, T. A. et al. Finding the missing heritability of complex diseases. Nature 461, 747–753 (2009).

6. Partridge, L. & Gems, D. Mechanisms of aging: public or private? Nat. Rev. Genet. 3, 165–175 (2002).

7. Benayoun, B. A., Pollina, E. A. & Brunet, A. Epigenetic regulation of ageing: linking environmental inputs to genomic stability. Nat. Rev. Mol. Cell Biol. 16, 593–610 (2015).

8. Di Francesco, A. et al. Dietary restriction impacts health and lifespan of genetically diverse mice. Nature 634, 684–692 (2024).

9. Litichevskiy, L. et al. Gut metagenomes reveal interactions between dietary restriction, ageing and the microbiome in genetically diverse mice. Nat. Microbiol. 10, 1240–1257 (2025).

10. Li, G.-H. et al. A multi-omics molecular landscape of 30 tissues in aging female rhesus macaques. Nat. Methods 22, 2658–2669 (2025).

11. Zhang, B. et al. Multi-omic rejuvenation and lifespan extension on exposure to youthful circulation. *Nat*. Aging 3, 948–964 (2023).

12. Zipple, M. N., Vogt, C. C. & Sheehan, M. J. Re-wilding model organisms: Opportunities to test causal mechanisms in social determinants of health and aging. Neurosci. Biobehav. Rev. 152, 105238 (2023).

13. Beura, L. K. et al. Normalizing the environment recapitulates adult human immune traits in laboratory mice. Nature 532, 512–516 (2016).

14. Ren, T. et al. Seasonal, spatial, and maternal effects on gut microbiome in wild red squirrels. Microbiome 5, 163 (2017).

15. Risely, A., Wilhelm, K., Clutton-Brock, T., Manser, M. B. & Sommer, S. Diurnal oscillations in gut bacterial load and composition eclipse seasonal and lifetime dynamics in wild meerkats. Nat. Commun. 12, 6017 (2021).

16. Stothart, M. R. et al. Methanogenic patterns in the gut microbiome are associated with survival in a population of feral horses. Nat. Commun. 15, 6012 (2024).

17. Reese, A. T. et al. Age Patterning in Wild Chimpanzee Gut Microbiota Diversity Reveals Differences from Humans in Early Life. Curr. Biol. 31, 613–620.e3 (2021).

18. Sadoughi, B., Schneider, D., Daniel, R., Schülke, O. & Ostner, J. Aging gut microbiota of wild macaques are equally diverse, less stable, but progressively personalized. Microbiome 10, 95 (2022).

19. Anderson, J. A. et al. High social status males experience accelerated epigenetic aging in wild baboons. eLife 10, e66128 (2021).

20. Snyder-Mackler, N. et al. Social determinants of health and survival in humans and other animals. Science 368, eaax9553 (2020).

21. Schumacher, A. E. et al. Global age-sex-specific all-cause mortality and life expectancy estimates for 204 countries and territories and 660 subnational locations, 1950–2023: a demographic analysis for the Global Burden of Disease Study 2023. The Lancet 406, 1731–1810 (2025).

22. Senior, A. M., Nakagawa, S., Raubenheimer, D. & Simpson, S. J. Global associations between macronutrient supply and age-specific mortality. Proc. Natl. Acad. Sci. 117, 30824–30835 (2020).

23. Crous-Bou, M. et al. Mediterranean diet and telomere length in Nurses’ Health Study: population based cohort study. BMJ 349, g6674 (2014).

24. Chen, C., Ding, S. & Wang, J. Digital health for aging populations. Nat. Med. 29, 1623–1630 (2023).

25. Nussey, D. H., Froy, H., Lemaitre, J.-F., Gaillard, J.-M. & Austad, S. N. Senescence in natural populations of animals: Widespread evidence and its implications for bio-gerontology. Ageing Res. Rev. 12, 214–225 (2013).

26. Kaeberlein, M., Creevy, K. E. & Promislow, D. E. L. The dog aging project: translational geroscience in companion animals. Mamm. Genome 27, 279–288 (2016).

27. Fleming, J. M., Creevy, K. E. & Promislow, D. E. L. Mortality in North American Dogs from 1984 to 2004: An Investigation into Age-, Size-, and Breed-Related Causes of Death. J. Vet. Intern. Med. 25, 187–198 (2011).

28. Kraus, C., Pavard, S. & Promislow, D. E. L. The Size–Life Span Trade-Off Decomposed: Why Large Dogs Die Young. Am. Nat. 181, 492–505 (2013).

29. Prescott, J. et al. Rationale and design of the Dog Aging Project precision cohort: a multi-omic resource for longitudinal research in geroscience. GeroScience 47, 5725–5748 (2025).

30. Creevy, K. E., Akey, J. M., Kaeberlein, M. & Promislow, D. E. L. An open science study of ageing in companion dogs. Nature 602, 51–57 (2022).

31. Bamberger, T. et al. Mapping the canine gut microbiome: insights from the Dog Aging Project. Nat. Commun. (2026) doi:10.1038/s41467-026-73193-y.

32. Harrison, B. R. et al. Protein Catabolites as Blood-Based Biomarkers of Aging Physiology: Findings From the Dog Aging Project. Aging Cell 24, e70226 (2025).

33. McCoy, B. M. et al. Aging at scale: Younger dogs and larger breeds from the Dog Aging Project show accelerated epigenetic aging. Preprint at 10.1101/2024.10.03.616519 (2024).

34. Mariner, B. L., et al. DNA methylation of transposons pattern aging differences across a diverse cohort of dogs from the Dog Aging Project. Preprint at 10.1101/2024.10.08.617286 (2025).

35. Roy, R. et al. DNA methylation signatures reveal that distinct combinations of transcription factors specify human immune cell epigenetic identity. Immunity 54, 2465–2480.e5 (2021).

36. Wikoff, W. R. et al. Metabolomics analysis reveals large effects of gut microflora on mammalian blood metabolites. Proc. Natl. Acad. Sci. 106, 3698–3703 (2009).

37. Dekkers, K. F. et al. An online atlas of human plasma metabolite signatures of gut microbiome composition. Nat. Commun. 13, 5370 (2022).

38. Argelaguet, R. et al. MOFA+: a statistical framework for comprehensive integration of multi-modal single-cell data. Genome Biol. 21, 111 (2020).

39. Horvath, S. & Raj, K. DNA methylation-based biomarkers and the epigenetic clock theory of ageing. Nat. Rev. Genet. 19, 371–384 (2018).

40. Muller, E., Shiryan, I. & Borenstein, E. Multi-omic integration of microbiome data for identifying disease-associated modules. Nat. Commun. 15, 2621 (2024).

41. Nikolich-Žugich, J. The twilight of immunity: emerging concepts in aging of the immune system. Nat. Immunol. 19, 10–19 (2018).

42. Goronzy, J. J. & Weyand, C. M. Mechanisms underlying T cell ageing. Nat. Rev. Immunol. 19, 573–583 (2019).

43. Sol, J. et al. Plasma acylcarnitines and gut-derived aromatic amino acids as sex-specific hub metabolites of the human aging metabolome. Aging Cell 22, e13821 (2023).

44. Panyard, D. J., Yu, B. & Snyder, M. P. The metabolomics of human aging: Advances, challenges, and opportunities. Sci. Adv. 8, eadd6155 (2022).

45. Kim, C.-S., Jung, S., Hwang, G.-S. & Shin, D.-M. Gut microbiota indole-3-propionic acid mediates neuroprotective effect of probiotic consumption in healthy elderly: A randomized, double-blind, placebo-controlled, multicenter trial and *in vitro* study. Clin. Nutr. 42, 1025–1033 (2023).

46. Yao, S. et al. Implications of metabolism on multi-systems healthy aging across the lifespan. Aging Cell 23, e14090 (2024).

47. Ghosh, T. S., Shanahan, F. & O’Toole, P. W. The gut microbiome as a modulator of healthy ageing. Nat. Rev. Gastroenterol. Hepatol. 19, 565–584 (2022).

48. Tett, A., Pasolli, E., Masetti, G., Ercolini, D. & Segata, N. Prevotella diversity, niches and interactions with the human host. Nat. Rev. Microbiol. 19, 585–599 (2021).

49. Turcsán, B. & Kubinyi, E. Differential behavioral aging trajectories according to body size, expected lifespan, and head shape in dogs. GeroScience 46, 1731–1754 (2024).

50. Harrison, B. R. et al. Dogs and humans share biomarkers of mortality. J. Gerontol. Ser. A 81, glaf279 (2026).

51. Huang, J. et al. Serum Metabolomic Profiling of All-Cause Mortality: A Prospective Analysis in the Alpha-Tocopherol, Beta-Carotene Cancer Prevention (ATBC) Study Cohort. Am. J. Epidemiol. 187, 1721–1732 (2018).

52. Mardinoglu, A. et al. The gut microbiota modulates host amino acid and glutathione metabolism in mice. Mol. Syst. Biol. 11, MSB156487 (2015).

53. Lehallier, B. et al. Undulating changes in human plasma proteome profiles across the lifespan. Nat. Med. 25, 1843–1850 (2019).

54. Shen, X. et al. Nonlinear dynamics of multi-omics profiles during human aging. *Nat*. Aging 4, 1619–1634 (2024).

55. Kirkwood, T. B. L. Deciphering death: a commentary on Gompertz (1825) ‘On the nature of the function expressive of the law of human mortality, and on a new mode of determining the value of life contingencies’. Philos. Trans. R. Soc. B Biol. Sci. 370, 20140379 (2015).

56. Wilmanski, T. et al. Blood metabolome predicts gut microbiome α-diversity in humans. Nat. Biotechnol. 37, 1217–1228 (2019).

57. Jaffe, A. E. & Irizarry, R. A. Accounting for cellular heterogeneity is critical in epigenome-wide association studies. Genome Biol. 15, R31 (2014).

58. Bergstedt, J. et al. The immune factors driving DNA methylation variation in human blood. Nat. Commun. 13, 5895 (2022).

59. Zünd, J. N. et al. Novel cross-feeding human gut microbes metabolizing tryptophan to indole-3-propionate. Gut Microbes 17, 2501195 (2025).

60. Saha, P. P. et al. Gut microbe-generated phenylacetylglutamine is an endogenous allosteric modulator of β2-adrenergic receptors. Nat. Commun. 15, 6696 (2024).

61. Brydges, C. R. et al. Indoxyl sulfate, a gut microbiome-derived uremic toxin, is associated with psychic anxiety and its functional magnetic resonance imaging-based neurologic signature. Sci. Rep. 11, 21011 (2021).

62. Chen, Q., et al. OMICmAge: An integrative multi-omics approach to quantify biological age with electronic medical records. bioRxiv 2023.10.16.562114 (2023) doi:10.1101/2023.10.16.562114.

63. Macdonald-Dunlop, E. et al. A catalogue of omics biological ageing clocks reveals substantial commonality and associations with disease risk. Aging 14, 623–659 (2022).

64. Shen, X. et al. Nonlinear dynamics of multi-omics profiles during human aging. *Nat*. Aging 4, 1619–1634 (2024).

65. Seshadri, G. et al. Immune cells are associated with mortality: the Health and Retirement Study. Front. Immunol. 14, 1280144 (2023).

66. Yang, J. et al. Development and validation of a blood biomarker score for predicting mortality risk in the general population. J. Transl. Med. 21, 471 (2023).

67. Batty, G. D., Gale, C. R., Kivimäki, M., Deary, I. J. & Bell, S. Comparison of risk factor associations in UK Biobank against representative, general population based studies with conventional response rates: prospective cohort study and individual participant meta-analysis. BMJ 368, m131 (2020).

68. Chen, F. L. et al. Evaluating instruments for assessing healthspan: a multi-center cross-sectional study on health-related quality of life (HRQL) and frailty in the companion dog. GeroScience 45, 2089–2108 (2023).

69. McCoy, B. M. et al. Social determinants of health and disease in companion dogs: a cohort study from the Dog Aging Project. Evol. Med. Public Health 11, 187–201 (2023).

70. Nyholm, L., et al. Holo-Omics: Integrated Host-Microbiota Multi-omics for Basic and Applied Biological Research. iScience 23, (2020).

71. Gomez-Cabrero, D. et al. Data integration in the era of omics: current and future challenges. BMC Syst. Biol. 8, I1 (2014).

72. Van Gassen, S., Gaudilliere, B., Angst, M. S., Saeys, Y. & Aghaeepour, N. CytoNorm: A Normalization Algorithm for Cytometry Data. Cytometry A 97, 268–278 (2020).

73. Kirwan, J. A., Weber, R. J. M., Broadhurst, D. I. & Viant, M. R. Direct infusion mass spectrometry metabolomics dataset: a benchmark for data processing and quality control. Sci. Data 1, 140012 (2014).

74. Altmann, A., Toloşi, L., Sander, O. & Lengauer, T. Permutation importance: a corrected feature importance measure. Bioinformatics 26, 1340–1347 (2010).

75. Dixon, P. VEGAN, a package of R functions for community ecology. J. Veg. Sci. 14, 927–930 (2003).

76. Davies, R. B. Hypothesis testing when a nuisance parameter is present only under the alternative. Biometrika 74, 33–43 (1987).

77. Dorrestein, P. C., Mazmanian, S. K. & Knight, R. Finding the Missing Links among Metabolites, Microbes, and the Host. Immunity 40, 824–832 (2014).

78. Sharon, G. et al. Specialized Metabolites from the Microbiome in Health and Disease. Cell Metab. 20, 719–730 (2014).

79. Zünd, J. N. et al. Novel cross-feeding human gut microbes metabolizing tryptophan to indole-3-propionate. Gut Microbes 17, 2501195 (2025).

80. Kuo, S. Z. et al. Associations between urinary 3-indoxyl sulfate, a gut microbiome-derived biomarker, and patient outcomes after intensive care unit admission. J. Crit. Care 63, 15–21 (2021).

81. Zhao, Z.-H. et al. Indole-3-propionic acid inhibits gut dysbiosis and endotoxin leakage to attenuate steatohepatitis in rats. Exp. Mol. Med. 51, 1–14 (2019).

82. Huang, J.-K. et al. Identification of Gut Microbiome Signatures Associated with Indole Pathway in Tryptophan Metabolism in Patients Undergoing Hemodialysis. Biomolecules 14, 623 (2024).

83. Koestler, D. C. et al. Blood-based profiles of DNA methylation predict the underlying distribution of cell types. Epigenetics 8, 816–826 (2013).

84. Houseman, E. A. et al. DNA methylation arrays as surrogate measures of cell mixture distribution. BMC Bioinformatics 13, 86 (2012).

85. Meryk, A. et al. Fcμ receptor as a Costimulatory Molecule for T Cells. Cell Rep. 26, 2681–2691.e5 (2019).

86. Kijas, J. M. H. et al. A Missense Mutation in the β-2 Integrin Gene (*ITGB2*) Causes Canine Leukocyte Adhesion Deficiency. Genomics 61, 101–107 (1999).

