## Supplementary Text for "Multi-omic longitudinal profiling of companion dogs reveals cross-omic coordination and structured aging trajectories"

---

### Supplementary Text

##### Microbiome-based prediction of circulating metabolites and comparison with human microbiome–metabolome associations

Given the strong global microbiome-metabolome association observed in the main analysis, as well as prior observations in human studies that link gut microbial composition to fecal and circulating metabolites[77,78], we set out to examine the relationship between the microbiome and metabolome profiles in our data more closely. Specifically, we trained predictive models using species-level relative abundances to predict metabolite levels and evaluated out-of-sample performance to identify metabolites that can be well predicted by the composition of the microbiome and thus potentially governed by microbial metabolism. Seven metabolites were found to be well predicted (Spearman's $r > 0.3$, $FDR < 0.1$; Fig. 2B-C, Supplementary Fig. 2 and Supplementary Table 1): indole-3-propionate ($r = 0.57$), 3-indoxyl sulfate ($r = 0.47$), glutaric acid ($r = 0.43$), phenylacetylglutamine ($r = 0.34$), trigonelline ($r = 0.32$), creatine ($r = 0.31$), and imidazole propionate ($r = 0.30$).

To place these microbiome-predictable metabolites in biological context, we examined whether they had known links to microbial metabolism and found that several of these metabolites are indeed known to be influenced by microbial metabolism. For example, 3-indoxyl sulfate (3-IS) and indole-3-propionate (IPA), which were the most accurately predicted metabolites, are both derivatives of indole and are produced downstream of bacterial tryptophan catabolism[36,79]. Specifically, 3-IS is generated when gut microbes convert tryptophan to indole, which is then sulfated by the host and excreted in urine (with reduced urinary 3-IS previously proposed as a marker in critical illness[80]). IPA, in turn, exhibits anti-inflammatory and gut-protective properties, promotes barrier integrity, and shows beneficial metabolic and neurological effects in animal models[81]. Finally, phenylacetylglutamine arises from microbial metabolism of dietary phenylalanine and has been implicated in cardiovascular disease via adrenergic receptor signaling[60]. Together, these results support the ability of the predictive models to recover metabolites with known or plausible links to microbial metabolism.

We further calculated permutation-based feature-importance profiles (using the Altmann method) for the models of the seven well-predicted metabolites to identify taxa contributing to prediction, retaining microbial species whose feature-importance p-values were significant after multiple-test correction (FDR < 0.1; Supplementary Fig. 2C and Supplementary Table 2). Across these metabolites, 210 unique microbial species showed significant importance, with a mean of 50.00 ± 27.18 (mean ± s.d.) contributing species per metabolite. The number of contributing species varied across metabolites, with higher counts for indole-3-propionate (91), 3-indoxyl sulfate (71), and creatine (70), and fewer contributors for glutaric acid (38), imidazole propionate (36), trigonelline (23) and phenylacetylglutamine (21). Several taxa were associated with the prediction of more than one metabolite. For example, multiple Prevotella species contributed to both indole-3-propionate and creatine prediction, and two Fusobacterium species contributed to both indole-3-propionate and 3-indoxyl sulfate. For 3-indoxyl sulfate, contributing taxa were primarily assigned to the genera Fusobacterium, Bacteroides, and Blautia. While prior work has implicated Bacteroides in indole metabolism[82], we did not observe direct evidence linking Fusobacterium or Blautia to 3-indoxyl sulfate production in our data, and their importance may reflect indirect or compositional associations. Overall, these results suggest that the microbiome-based prediction of circulating metabolites is not driven by a single dominant taxon but rather reflects broader microbial community structure and potentially shared metabolic pathways across taxa.

Finally, to assess whether metabolites that were well predicted based on the microbiome in our cohort exhibit similar microbiome-related patterns in humans, we compared our predictability estimates with microbiome-metabolite associations reported in a population-based Swedish CardioPulmonary bioImage Study (SCAPIS; n = 8,583)[37]. Examining the metabolites that were measured in both datasets, we found a significant correlation between our microbiome-based predictability values and the variance explained by the microbiome in this human serum cohort (Spearman $r = 0.239$, $p < 0.05$; Fig. 2D; Supplementary Table 3), with several well-predicted metabolites in our cohort ranked among metabolites with the highest microbiome-attributable variance in the human dataset. This suggests partial conservation of microbiome-metabolome associations across species.

**Methylation-based prediction of immune-cell composition**

Motivated by the strong global association between methylation and FACS profiles observed in the main analysis, and consistent with the known sensitivity of blood methylation profiles to immune-cell composition[83,84], we examined whether PBMC methylation profiles capture variation in immune-cell composition measured by flow cytometry. Using machine-learning models trained to predict each of the 60 FACS-derived immune-cell features from methylation data, we identified 15 immune-cell features that were well predicted (Spearman's $r > 0.3$, $FDR < 0.1$; Supplementary Fig. 3A,B and Supplementary Table 4). Prediction performance among these features ranged from $r = 0.31$ to $r = 0.49$. The most accurately predicted features were the median side scatter of double-negative cells (DN_med_SS; $r = 0.49$), the median side scatter of CD8+ cells (CD8_med_SS; $r = 0.49$), and the side scatter of CD5-high cells (CD5hi_SS; $r = 0.48$).

To identify the epigenomic features contributing to these predictions, we examined promoter-level feature importance across the FACS prediction models. Of 3,499 promoter-associated methylation features, 350 showed significant positive predictive importance ($FDR < 0.1$; Supplementary Fig. 3C) for at least one FACS measure, comprising 672 significant promoter-FACS associations. Many promoters contributed to multiple immune-cell measurements: 116 were significantly informative for at least two FACS measures, 61 for at least three, and 21 for at least five (Supplementary Fig. 3D). Several recurrently informative promoters mapped to genes with established immune functions; FCMR (Fc mu receptor) was particularly prominent, ranking among the top 1% of predictors for 23 of the 60 FACS measures and showing significant predictive importance for 14. ITGB2 (integrin subunit beta 2; CD18) ranked among the top 1% for 10 measures and was significantly informative for seven. These genes have well-established roles in lymphocyte biology and leukocyte adhesion and trafficking, respectively, providing biologically interpretable examples of the epigenomic features underlying the methylation-based predictions[85,86].

PBMC methylation profiles reflect the combined methylation patterns of multiple immune-cell populations, and these results therefore likely capture variation in underlying immune-cell composition and immune-state variation.

### Supplementary Figures

#### Supplementary Figure 1

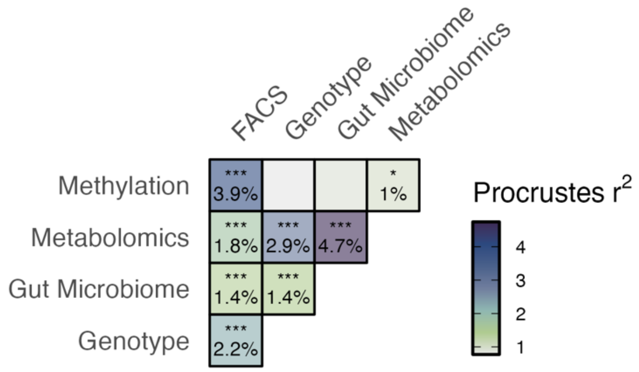


**Supplementary Figure 1.** Pairwise Procrustes analysis quantifying concordance between omic domains; tiles are colored by the Procrustes correlation (higher = stronger concordance). Asterisks denote FDR-adjusted significance (· FDR ≤ 0.1; * ≤ 0.05; ** ≤ 0.01; *** ≤ 0.001).

#### Supplementary Figure 2

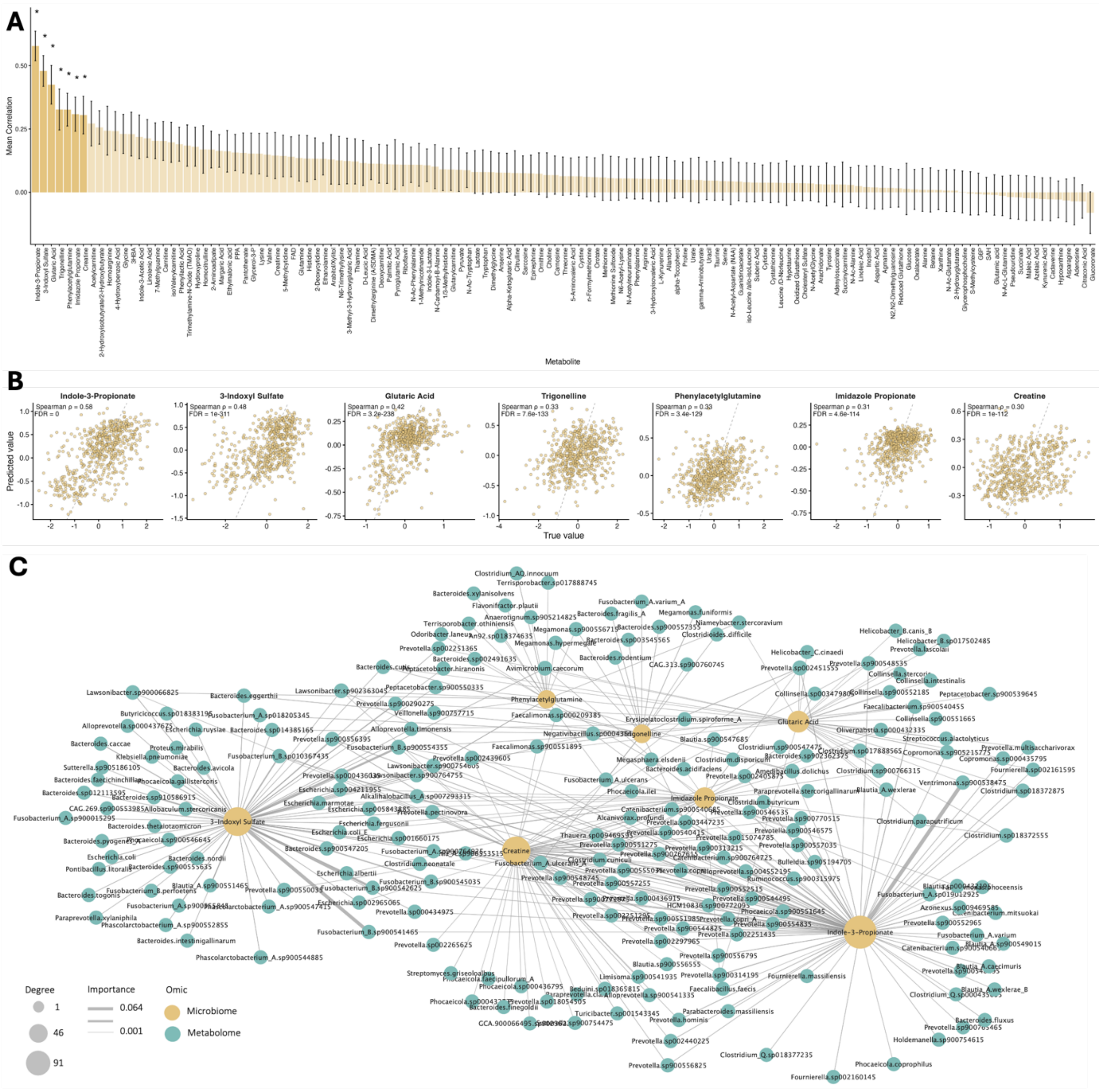


**Supplementary Figure 2. Microbiome-metabolome associations. (A)** Predictability of metabolites: metabolites ranked by cross-validated correlation (|Spearman r|) between observed and predicted concentrations. Well predicted metabolites (Absolute Spearman r between real and predicted values, FDR < 0.1) are marked with an asterisk and shown in saturated color. **(B)** Observed vs. predicted values for the well predicted metabolites. **(C)** Feature-importance network for the seven well-predicted metabolites. Nodes represent metabolites (yellow) and microbial species (light blue); edges link species to metabolites for which they show significant Altmann permutation importance (FDR < 0.1). Edge width encodes importance magnitude; node size encodes degree.

#### Supplementary Figure 3

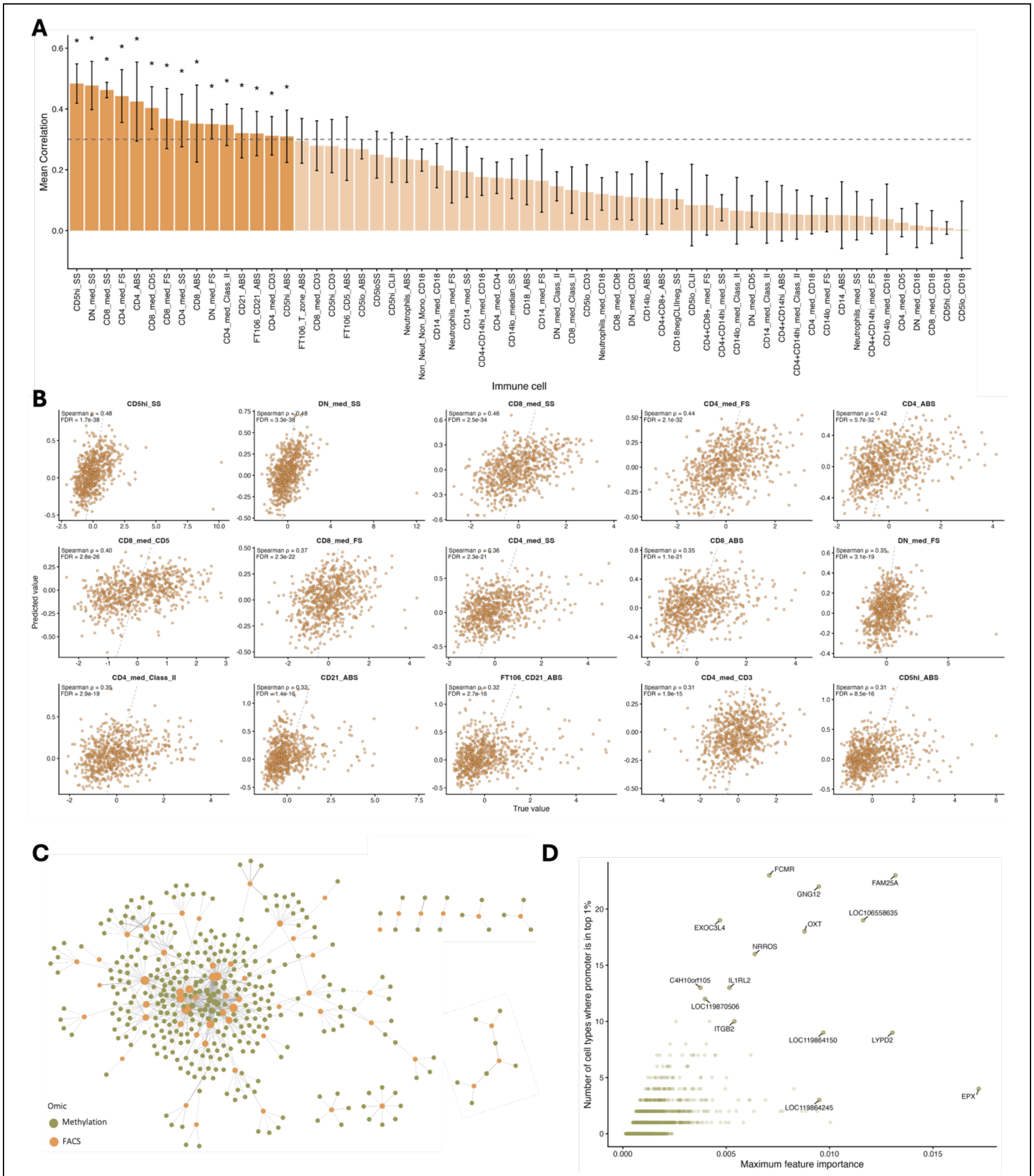


**Supplementary Figure 3. Methylation-FACS associations. (A-C)** Similar to Supplementary Figure 2A-C, but using methylation data to predict immune cell feature values. **(D)** Promoter-level feature importance across FACS prediction models. Each point represents a promoter-associated gene. The x-axis shows the maximum feature importance across the 60 FACS measures, and the y-axis shows the number of measures for which the promoter ranked among the top 1% of predictive features. The 15 genes with the highest combined score for maximum importance and recurrence across FACS measures are labeled.

#### Supplementary Figure 4

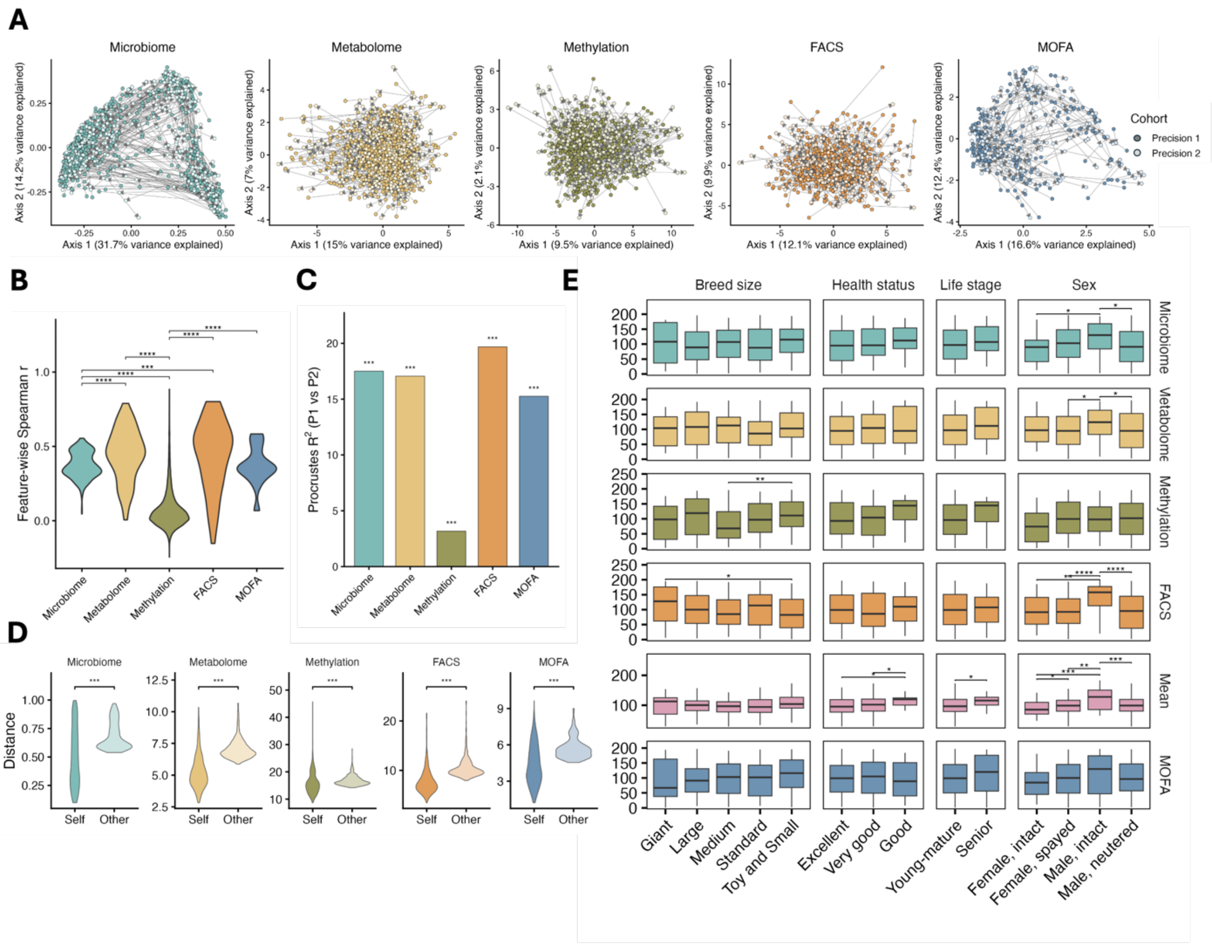


**Supplementary Figure 4. Longitudinal dynamics across omics. (A)** Principal Coordinate Analysis (PCoA) for each omic/MOFA+ latent factors, using Bray-Curtis distances for the microbiome and Euclidean distances for all other omics; points are color-coded by sampling year and by omic, and arrows connect each Precision 1 (P1) sample to its matched Precision 2 (P2) sample. **(B)** Violin plots showing feature-wise temporal stability per omic (Spearman correlation for each feature between P1 and P2 across dogs). Differences were tested with Wilcoxon tests. **(C)** Bar plot of Procrustes statistics quantifying global cross-time concordance (P1 vs. P2) for each omic. Significance was assessed with permutation tests. **(D)** Violin plots comparing each dog's P1-to-P2 self-distance with the mean distance from its P1 profile to the P2 profiles of all other dogs. Differences were tested using linear mixed-effects models adjusted for breed size, sex/sterilization class, and life stage, with a random intercept for dog to account for the two distance measurements contributed by each dog. **(E)** Boxplots of demographic correlates of longitudinal change: x-axis, demographic categories; y-axis, ranked P1 to P2 distance. Group differences were tested with Wilcoxon procedures. In panels B-E, p-values were FDR-adjusted and significance is denoted by asterisks: * FDR < 0.05, ** FDR < 0.01, *** FDR < 0.001, **** FDR < 0.0001. In all panels, colors denote omic.

#### Supplementary Figure 5

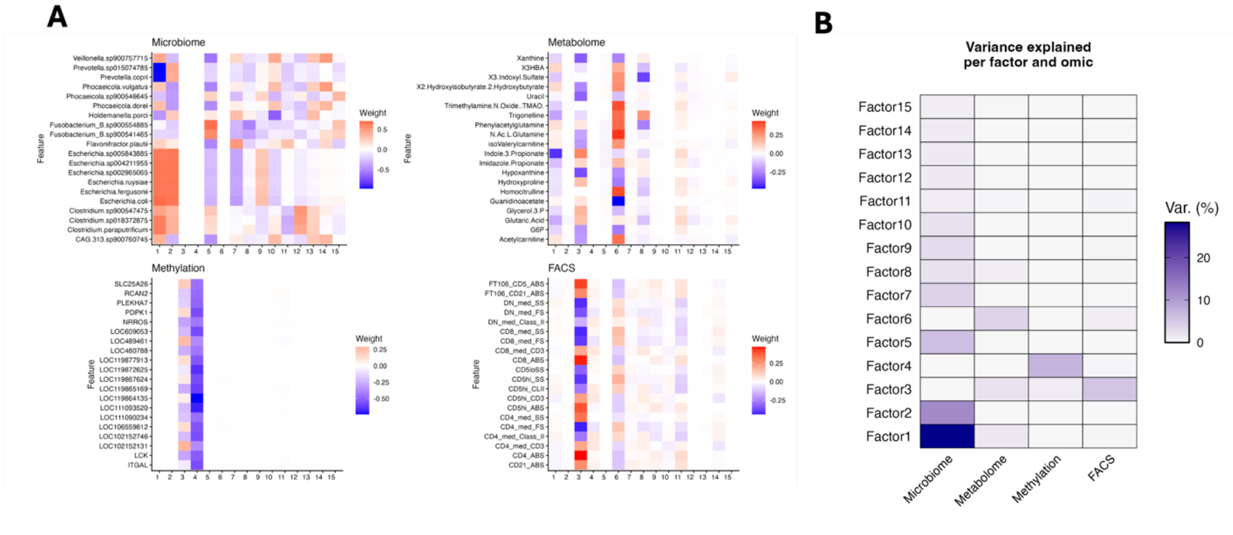


**Supplementary Figure 5. MOFA+ latent factor analysis**. **(A)** Heatmaps show the top 20 features per omic (ranked by total absolute weight across factors) for each omic. Columns are MOFA+ factors (shared ordering across panels); rows are features. Tile color indicates the factor loading (blue = negative, white = near zero, red = positive). **(B)** Heatmap shows the percent variance explained per factor (rows) in each omic view (columns). Only factors with ≥1% total variance are displayed. Color scale indicates explained variance (%).

#### Supplementary Figure 6

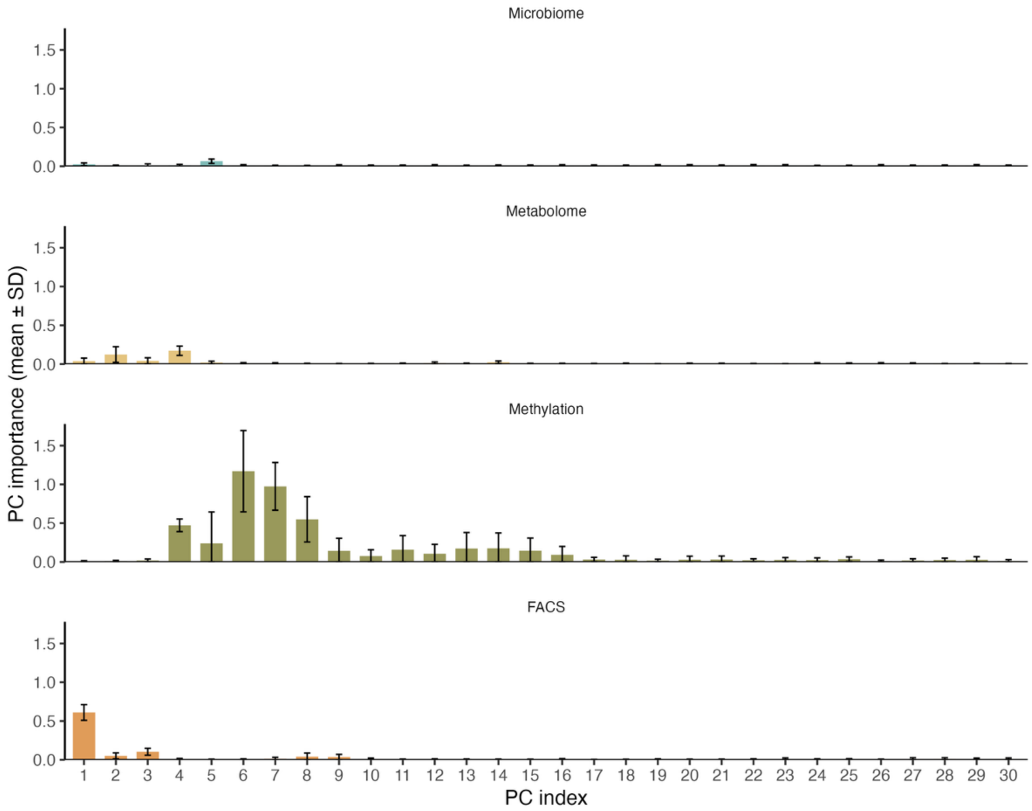


**Supplementary Figure 6. PC importance by omic for age prediction.** Altmann permutation-based PC importances grouped by omic; error bars show SD across folds and repeats.

#### Supplementary Figure 7

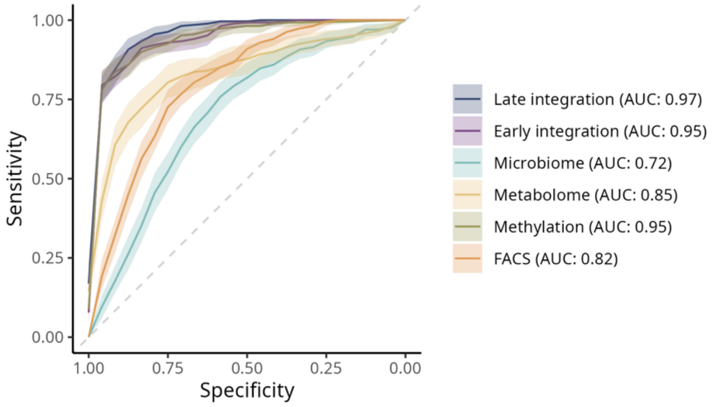


**Supplementary Figure 7. Age-category prediction including methylation data**. ROC curves for age-category, as in Figure 4D, but with methylation data included.

#### Supplementary Figure 8

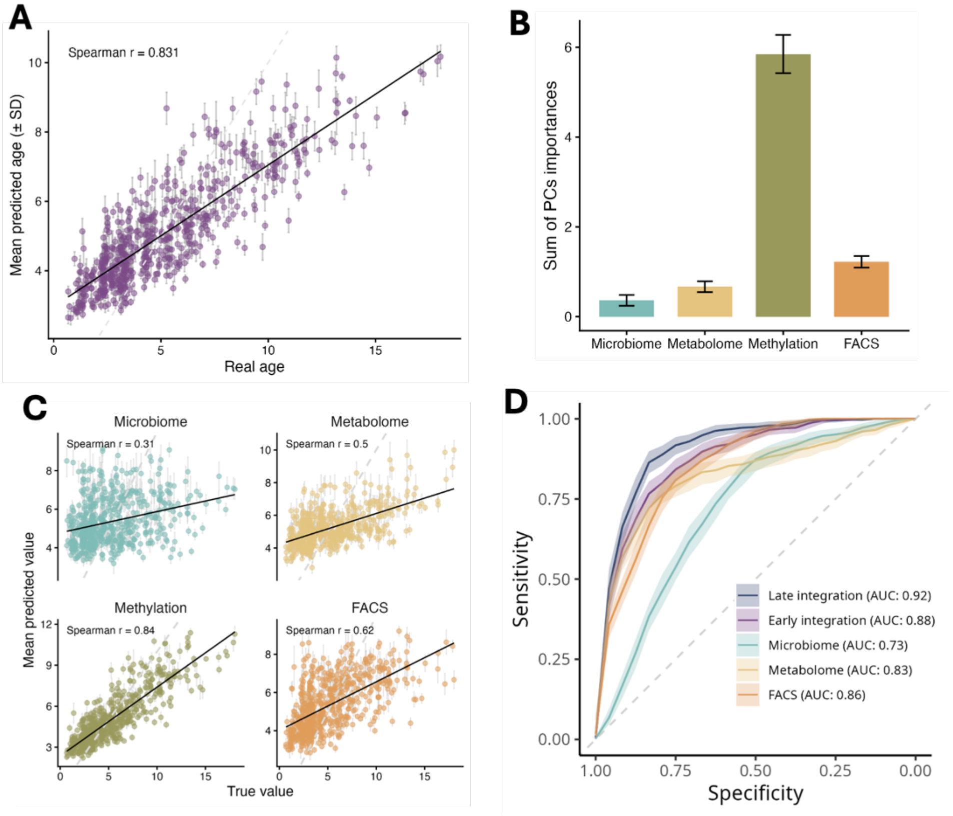


**Supplementary Figure 8. Association of multiple omic domains with age in the full cohort.** Same as in Figure 4A-D, but including all dogs, irrespective of health status.

**Supplementary Figure 9**

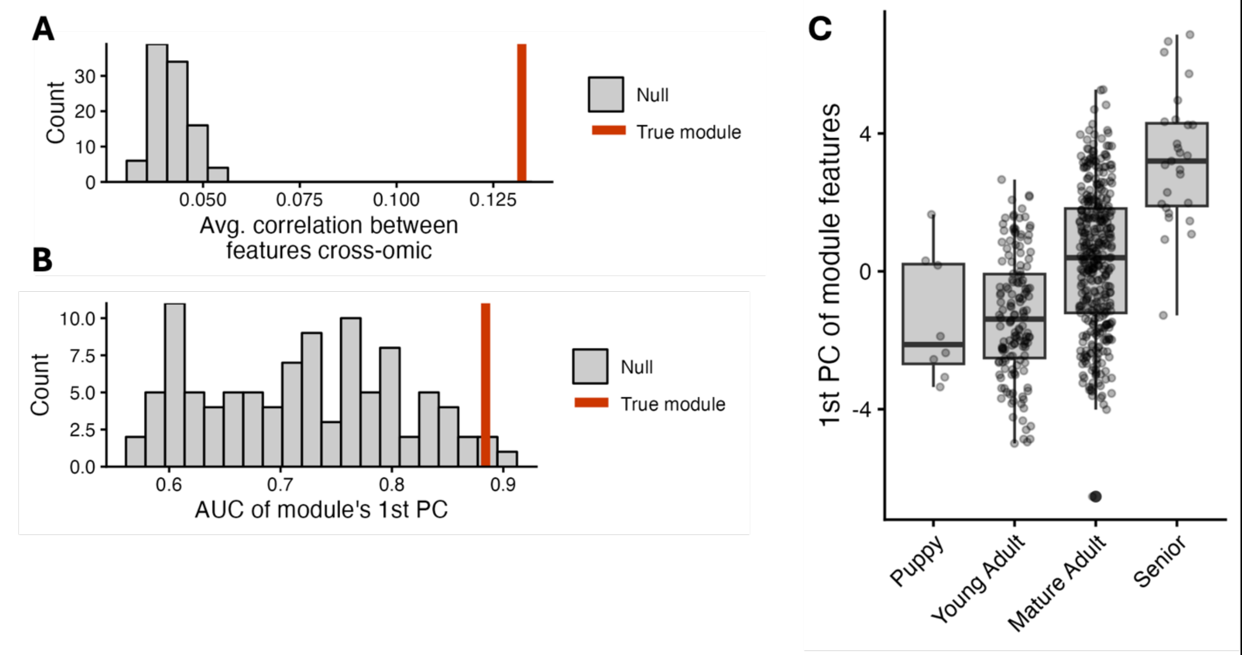


**Supplementary Figure 9. Evaluation of the MintTea module from Figure 4E. (A)** Mean cross-omic feature–feature correlation for the real module (red) versus size- and omic-matched null modules (gray). **(B)** Cross-validated AUROC of the module's first principal component (PC1) for age classification; red = real module, gray = nulls. **(C)** PC1 values across dog life stages for features in the module.

**Supplementary Figure 10**

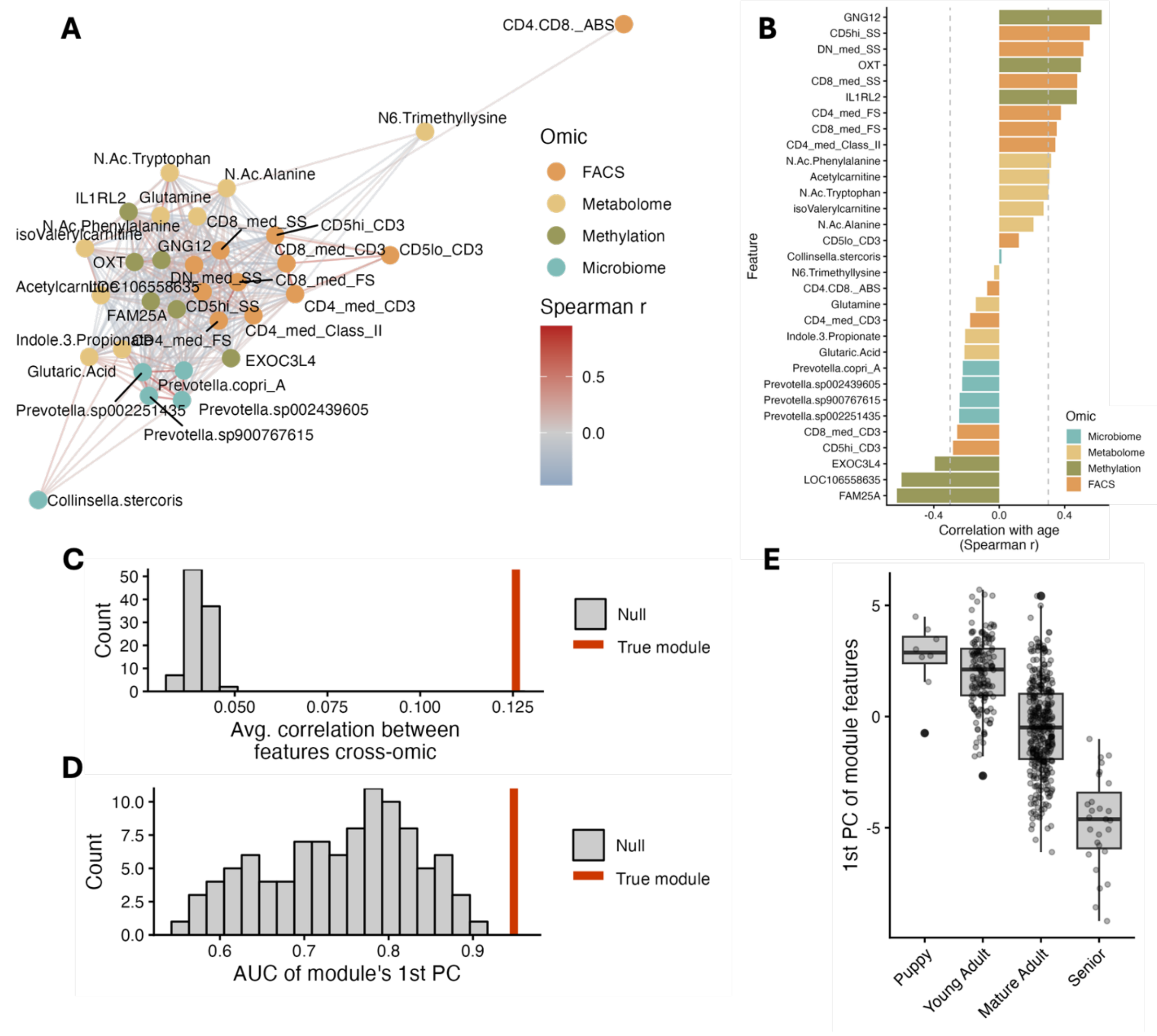


**Supplementary Figure 10. MintTea module including methylation data. (A-B)** MintTea module network and feature-age correlations, shown as in Figure 4E-F, but including methylation features. **(C-E)** Evaluation of the same module, as in Supplementary Figure 9A-C
